# Body mass index, time of day, and genetics affect perivascular spaces in the white matter

**DOI:** 10.1101/2020.06.20.162404

**Authors:** Giuseppe Barisano, Farshid Sepehrband, Nasim Sheikh-Bahaei, Meng Law, Arthur W. Toga

## Abstract

The analysis of cerebral perivascular spaces (PVS) using magnetic resonance imaging (MRI) allows to explore *in vivo* their contributions to neurological disorders. To date the normal amount and distribution of PVS in healthy human brains are not known, thus hampering our ability to define with confidence pathogenic alterations. Furthermore, it is unclear which biological factors can influence the presence and size of PVS on MRI. We performed exploratory data analysis of PVS volume and distribution in a large population of healthy individuals (n = 897, age = 28.8 ± 3.7). Here we describe the global and regional amount of PVS in the white matter, which can be used as a reference for clinicians and researchers investigating PVS and may help the interpretation of the structural changes affecting PVS in pathological states. We found a relatively high inter-subject variability in the PVS amount in this population of healthy adults (range: 1.31-14.49 cm^3^). We then identified body mass index, time of day, and genetics as new elements significantly affecting PVS *in vivo* under physiological conditions, offering a valuable foundation to future studies aimed at understanding the physiology of perivascular flow.

## Introduction

Perivascular spaces (PVS), also known as Virchow-Robin spaces, are tube-shaped structures that surround perforating arteries and small blood vessels in the brain parenchyma, including arterioles, venules, and capillaries(1). PVS is a major component of the brain clearance system and accommodates the influx of CSF to the cerebral parenchyma through the peri-arterial space and the efflux of interstitial fluid to the lymphatic system through the peri-venous space(2,3). Detecting pathological PVS changes is of high clinical significance because it provides mechanistic insight into disease pathology, aids in diagnosis, and can be used for disease monitoring, as PVS alterations may precede and be more reversible than demyelination and axonal loss in neurodegenerative disorders(4,5). However, the physiological profile of the PVS is not fully understood, limiting the ability to identify and recognize PVS abnormalities in neurological disorders, especially in subclinical phases of the disease.

In the past two decades, improvements in imaging and post-processing techniques as well as the more widespread use of ultra-high field MRI systems supported significantly enhanced evaluation of PVS, and increasing attention has been dedicated to PVS, their pathophysiological variations, and their potential role as a diagnostic biomarker(6–9). In fact, to date increased PVS visibility on human MRI studies has been found associated with ageing(10) and a number of pathologic conditions, such as neuropsychiatric and sleep disorders(11–15), multiple sclerosis(16,17), mild traumatic brain injury(18,19), Parkinson’s disease(20), post-traumatic epilepsy(19), myotonic dystrophy(21), systemic lupus erythematosus(22), cerebral small vessel disease(23–27), and cerebral amyloid-β pathologies, including Alzheimer’s disease (AD) and Scerebral amyloid angiopathy (CAA)(28–31). These findings suggest that a higher number of PVS visible on MRI might be an indicator of impaired brain health, although not specific for any single disease.

Despite the increased interest in the role of PVS within the scientific community, there are several unsolved controversies regarding *in vivo* PVS analysis using MRI. Resolving these issues is critical for the interpretation of the results derived from PVS studies(5,32). Some of the main problems include: 1) the definition of the enlarged PVS: traditionally, the increased number of detected PVS has been interpreted as an enlargement of PVS, but there is no agreement regarding the radiological definition of enlarged PVS, as there is no quantitative measure of PVS in healthy people; 2) the visual scoring used in most studies focus on basal ganglia and centrum semiovale, but the regional distribution of PVS in the white matter is unknown; 3) the role and effect of clinical and genetic factors on the physiological amount of PVS have not been thoroughly investigated.

In this study, we provide the first quantitative analysis of PVS performed using submilliter MRI in a large population of 897 healthy adults from the human connectome project(33). We describe the regional distribution and extent of PVS in the white matter of the human brain, which can be used by researchers and clinicians as a normative atlas of PVS or as a reference resource. The age range of participants was between 22 and 37 years old and was chosen to represent healthy adults beyond the age of major neurodevelopmental changes and before the onset of neurodegenerative alterations(33). We also investigated the relationship between PVS and multiple demographic, clinical, and genetic parameters in order to understand which factors may significantly influence the amount of PVS in healthy adults.

## Material and Methods

### Study Population

A total of 897 participants were identified from the Human Connectome Project study (S900 release)(33). According to how the Human Connectome Project study has been designed and performed, recruiting efforts were aimed at ensuring that participants broadly reflect the ethnic and racial composition of the U.S. population as represented in the 2000 decennial census(34). The goal was to recruit a pool of individuals that is generally representative of the population at large, in order to capture a wide range of variability in healthy individuals with respect to behavioral, ethnic, and socioeconomic diversity(34). The study protocol was approved by the Institutional Review Board at the University of Southern California (IRB# HS-19-00448) conforming with the World Medical Association Declaration of Helsinki. Written consent was obtained from all participants at the beginning of the first day of involvement in the project(34). Only healthy individuals were included in the study. Inclusion and exclusion criteria are listed in supplementary table 1.

### Clinical and behavioral data

Collected demographic and clinical data included: age, sex, height and weight with the corresponding BMI, blood pressure, years of education, hematocrit, glycated hemoglobin, and thyroid stimulating hormone in blood. Information about alcohol consumption and tobacco smoking was collected through the Semi-Structured Assessment for the Genetics of Alcoholism interview (SSAGA)(35). The NIH Toolbox (http://www.nihtoolbox.org) was used to assess the domains of cognition, emotion, motor function, and sensation(33). Additionally, each participant underwent the Mini-Mental State Examination(36). The Pittsburgh Sleep Quality Index was used to evaluate sleep quality and quantity(37): specifically, the sleep quality was computed as the sum of 7 analyzed components including subjective sleep quality, sleep latency, sleep duration, habitual sleep efficiency, sleep disturbances, use of sleeping medication, and daytime dysfunction; the sleep quantity was assessed by asking the participant what was the average number of hours of actual sleep per night in the past month, not counting time falling asleep and getting out bed.

### MRI methods and analysis

The preprocessed T1-weighted (TR 2400 ms, TE 2.14 ms, TI 1000 ms, FOV 224 x 224 mm) and T2-weighted (TR 3200 ms, TE 5.65 ms, FOV 224 x 224 mm) images of the Human Connectome Project(38), acquired at 0.7 mm^3^ resolution on a Siemens 3T Skyra scanner (Siemens Medical Solutions, Erlangen, Germany), were used for the PVS analysis. A total of 45 participants underwent a second MRI scan, with the same protocol and scanner, after an interval of 139 ± 69 days; these scans were used to assess the effect of sleep and time of day with the PVS. The preprocessing steps included: correction for gradient nonlinearity, readout, and bias field; alignment to AC-PC subject space; registration to MNI 152 space using the FNIRT function in FSL(39); generation of individual cortical, white matter, and pial surfaces and volumes using the FreeSurfer software(40) and the HCP pipelines(38).

### PVS analysis

For PVS quantification and mapping, we first enhanced the visibility of PVS and then automatically segmented PVS across the white matter. We combined T1- and T2-weighted images that were adaptively filtered to remove non-structured high-frequency spatial noise by using a filtering patch which removes the noise at a single-voxel level and preserves signal intensities that are spatially repeated, thus preserving PVS voxels(41,42). Non-local mean was used for removing high frequency noise, which measures the image intensity similarities by considering the neighboring voxels in a blockwise fashion, where filtered image is 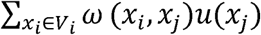. For each voxel (*x*_*j*_) the weight (*ω*) is measured using the Euclidean distance between 3D patches. The adaptive non-local mean filtering technique adds a regularization term to the above formulation to remove bias intensity of the Rician noise observed in MRI.

We then used n-tissue parcellation technique of the Advanced Normalization Tools (ANTs) package(43,44). Parcellated white matter was used as a mask for PVS analysis. For PVS segmentation, we first applied Frangi filter(45), using Quantitative Imaging Toolkit(46), which extracts the likelihood of a voxel belonging to a PVS. The Frangi filter has been shown to be an adequate tool for PVS segmentation(41,47–51). Frangi filter estimates a vesselness measure for each voxel from eigenvectors of the Hessian matrix of the image. Default parameters of *α* = 0.5, *β* = 0.5 and *c* were used, as recommended in (45). The parameter *c* was set to half the value of the maximum Hessian norm. Frangi filter estimated vesselness measures at different scales and provided the maximum likeliness. The scale was set to a large range of 0.1 to 5 voxels in order to maximize the vessel inclusion. The output of this step is a quantitative maximum likelihood map of vessels in regions of interest(45). We selected a previously optimized scaled threshold of 1.5 (equal to raw threshold of 1e-6) in the vessel map in order to obtain a binary mask of PVS regions, which is required for obtaining PVS volumetric measurements and spatial distribution(41).

The periventricular voxels were excluded via a dilated mask of the lateral ventricles in order to remove the incorrectly segmented PVS at the lateral ventricles-white matter boundary(41). This PVS segmentation technique has been previously validated on the same MRI dataset(41). For the optimization of the protocol, we visually analyzed the PVS masks of 100/897 MRI scans randomly selected. There was a good concordance between the measure obtained via our technique and the manual analysis of PVS independently performed by two expert PVS readers(41). Finally, PVS ratio was extracted across the white matter regions, parcellated based on Desikan-Killiany atlas using *FreeSurfer* software(52). The total PVS-white matter ratio was also estimated.

### Genome-Wide Association Analysis

Genome-wide single nucleotide polymorphisms (SNPs) genotyping was performed in 831/897 participants with useable blood or saliva-based genetic material. For this study, we used only samples processed with one custom microarray chip consisting of the Illumina Mega Chip (2 million multiethnic SNPs). Clinical and demographic data, PVS measurements, and SNPs were combined to yield a single data set for every individual. Genotype information was available for 2,119,803 typed SNPs across 831 individuals with clinical data and PVS measurements. The high number of SNPs available allowed us to perform a stringent SNP-level filtering: we filtered out SNPs for which the minor allele frequency was less than 1%, in order to ensure adequate power to infer a statistically significant relationship between the SNP and the PVS. After this step, 471,068 typed SNPs across 831 individuals persisted and underwent further pre-processing steps. We subsequently performed a sample-level filtering: a call rate of 100% was applied in order to include only participants’ samples with 100% of genetic data available; additionally, we excluded samples exhibiting deviations from the Hardy-Weinberg equilibrium with an inbreeding coefficient higher than 0.1, since excess heterozygosity across typed SNPs within an individual may be an indication of poor sample quality(53). For ancestry filtering, we first applied linkage disequilibrium pruning using a threshold value of 0.2, which eliminates a large degree of redundancy in the data and reduces the influence of chromosomal artifacts(54,55). Then, we used the Method of Moments procedure to calculate the identity by descent (IBD) kinship coefficient: pairwise IBD distances were computed to search for sample relatedness and participants with the highest number of pairwise kinship coefficients >0.1, which typically suggest relatedness, duplicates, or sample mixture, were iteratively removed(55). This resulted in the exclusion of 455 samples. Among non-twin and twin siblings included in this study, all but one member of each biologically independent sibship was filtered out at this step. At the end of the pre-processing procedure, 471,068 typed SNPs across 376 individuals were considered in the final genome-wide association analysis. A Bonferonni-corrected genome-wide significance threshold of 5×10^−8^ and a suggestive association significance threshold of 5×10^−6^ were adopted(55).

### Statistical Analysis

The statistical analysis was done using the R package version 1.2.5 (R Development Core Team, 2019).

The Shapiro-Wilk test for normality was used to assess data distribution. All data analyzed exhibited a distribution that was significantly different from normal distribution. Therefore, in the following non-parametric tests were applied: the Wilcoxon matched-pairs signed rank test was used to compare differences across paired groups, while the Wilcoxon rank sum (Mann-Whitney) test was performed to compare two unmatched groups. The Kruskal-Wallis test was used to compare three or more unmatched groups. Correlations were measured using the Spearman’s coefficient.

In order to assess which demographic and clinical parameters influenced the amount of PVS measured in the brain, general linear models were applied, using one clinical factor at a time as independent variable, and the PVS ratio as dependent variable. After the identification of potentially significant factors, we performed a new general linear model analysis including all of them together as independent variables and the PVS ratio as the dependent variable. The two-way ANCOVA model was used to test the effect of gender and BMI on the PVS ratio controlling for age.

When analyzing the relationship between PVS ratio and the results of the behavioral tests, a principal component analysis was initially applied to convert and reduce this set of variables into a set of linearly uncorrelated variables, since many of the behavioral scores were expected to have multi-collinearity. The first principal component, explaining most of the variance in behavioral measures, was then used to identify the most influential neurocognitive scores, which were employed in a series of linear models as dependent variables to investigate whether the PVS ratio is a predictor of cognitive performance. Regression models were fitted using the ordinary least square technique. The Benjamini-Hochberg method was adopted to correct for multiple comparisons with a false discovery rate of 0.05. All p-values were 2-sided and considered significant at <0.05.

## Results

### Analysis of PVS volume, ratio, and distribution across white matter regions

We were able to compute PVS volume in 897 participants (demographic and clinical data are reported in table 1).

**Table 1.** Demographic and clinical characteristics of participants from the Human Connectome Project (S900 Release) included in this study. N=897 unless otherwise specified. Data are mean ± standard deviation.

|  | Population |  |  |
| --- | --- | --- | --- |
|  | Male | Female | Overall |
| Age | 28 ± 3.65 | 29.46 ± 3.58 | 28.82 ± 3.68 |
| BMI, kg/m<sup>2</sup> (n=896) | 26.95 ± 4.52 | 26.42 ± 5.84 | 26.65 ± 5.3 |
| Education (years) | 14.77 ± 1.79 | 14.98 ± 1.84 | 14.89 ± 1.82 |
| Mini-mental state examination score (out of 30) | 28.96 ± 1.1 | 29.05 ± 0.96 | 29.01 ± 1.03 |
| Smoking history (n=896) |  |  |  |
| - Non-smokers | 186 | 304 | 490 |
| - Occasional smokers | 87 | 89 | 176 |
| - Regular smokers | 121 | 109 | 230 |
| Average number of cigarettes/day | 10.08 ± 5.66 | 9.72 ± 6.41 | 9.91 ± 6.02 |
| Average alcoholic drinks/week (n=880) | 6.47 ± 8.29 | 3.06 ± 4.34 | 4.54 ± 6.58 |
| Systolic Blood pressure (mmHg) (n=884) | 129.29 ± 13.6 | 120.13 ± 13.66 | 124.17 ± 14.36 |
| Diastolic Blood pressure (mmHg) (n=884) | 79.11 ± 10.34 | 75.22 ± 10.38 | 76.94 ± 10.54 |
| Hematocrit (%) (n=823) | 46.03 ± 3.57 | 40.87 ± 4.43 | 43.12 ± 4.81 |
| Blood Thyroid Hormone (mU/L) (n=610) | 1.73 ± 0.91 | 1.82 ± 1.31 | 1.78 ± 1.13 |
| Glycated Hemoglobin (%) (n=603) | 5.26 ± 0.41 | 5.26 ± 0.39 | 5.26 ± 0.4 |
| Average amount of sleep hours per night | 6.81 ± 1.15 | 6.82 ± 1.14 | 6.81 ± 1.15 |
| **Neuroimaging data** |  |  |  |
| Total intracranial volume (cm<sup>3</sup>) | 1702.86 ± 148.21 | 1470.94 ± 149.79 | 1572.81 ± 188.33 |
| Whole brain volume (cm<sup>3</sup>) | 1264.98 ± 103.12 | 1106.78 ± 85.79 | 1176.28 ± 122.31 |
| White matter volume (cm<sup>3</sup>) | 480.28 ± 50.41 | 415.27 ± 40.70 | 443.83 ± 55.54 |
| PVS Volume (mm<sup>3</sup>) | 5842.93 ± 2356.69 | 4393.76 ± 1733.64 | 5029.39 ± 2153.15 |
| PVS Ratio (%) | 1.22 ± 0.45 | 1.07 ± 0.39 | 1.14 ± 0.43 |

The mean PVS volume in the white matter was 5.03±2.15 cm^3^, with a high inter-subject variability (range: 1.31-14.49 cm^3^) (Figure 1).

**Figure 1.**
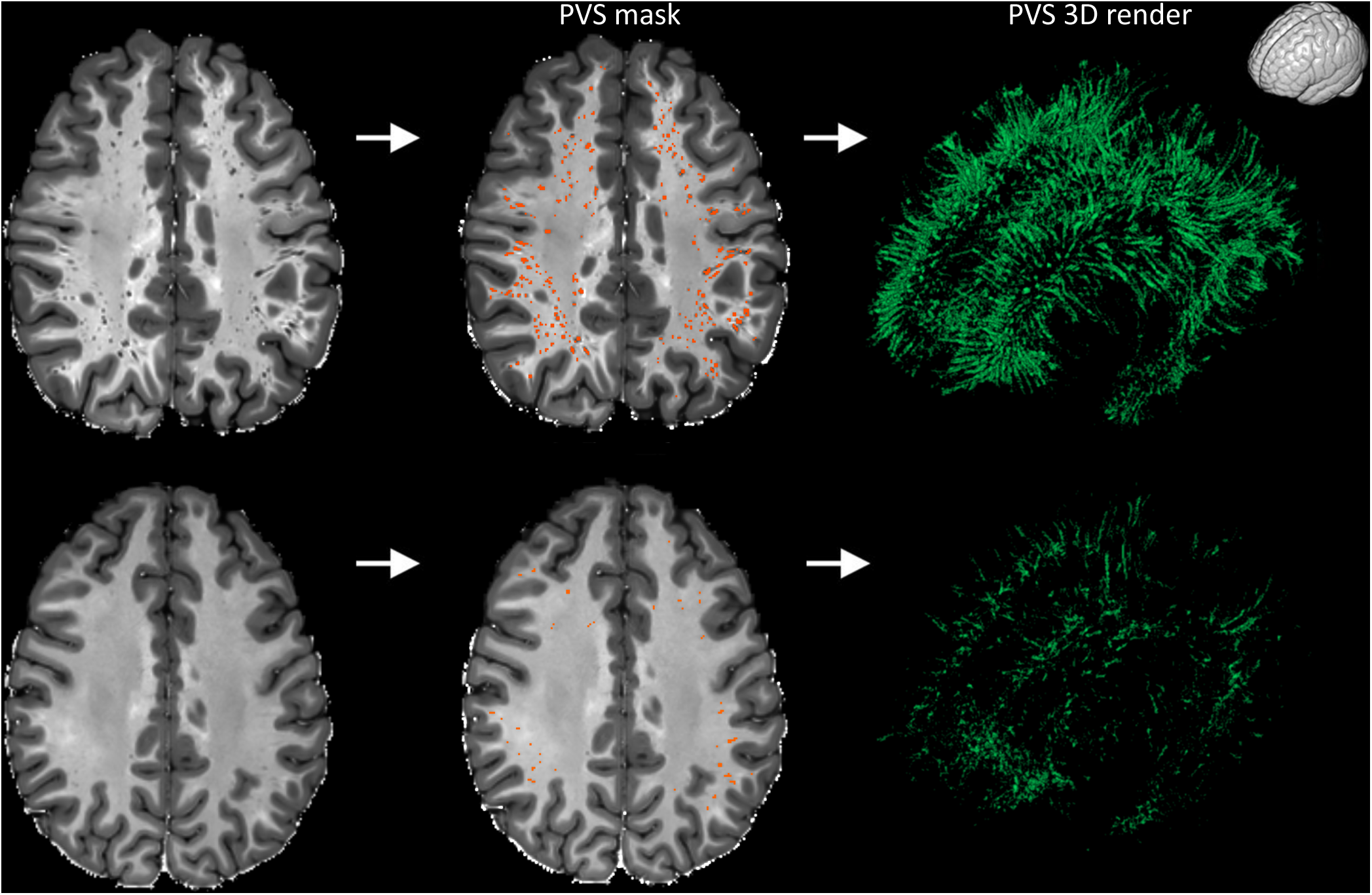
Examples showing the high inter-subject variability of perivascular spaces (PVS) in healthy participants. The participant on the top is a 27 years old female, while the participant on the bottom is a 32 years old male (two extreme cases are intentionally presented to highlight the high inter-subject variability in PVS). The MRI scans are shown on the left column and the PVS mask were overlaid in the center (orange). The images on the right are the corresponding 3D maps of the PVS masks. The orientation of the 3D maps is reported on the top right corner.

Among the regions of interest (ROIs) segmented in the white matter, the superior frontal and parietal regions showed the highest percentage of PVS, including on average more than 8% and 6% of the total PVS volume, respectively (Figure 2A).

**Figure 2.**
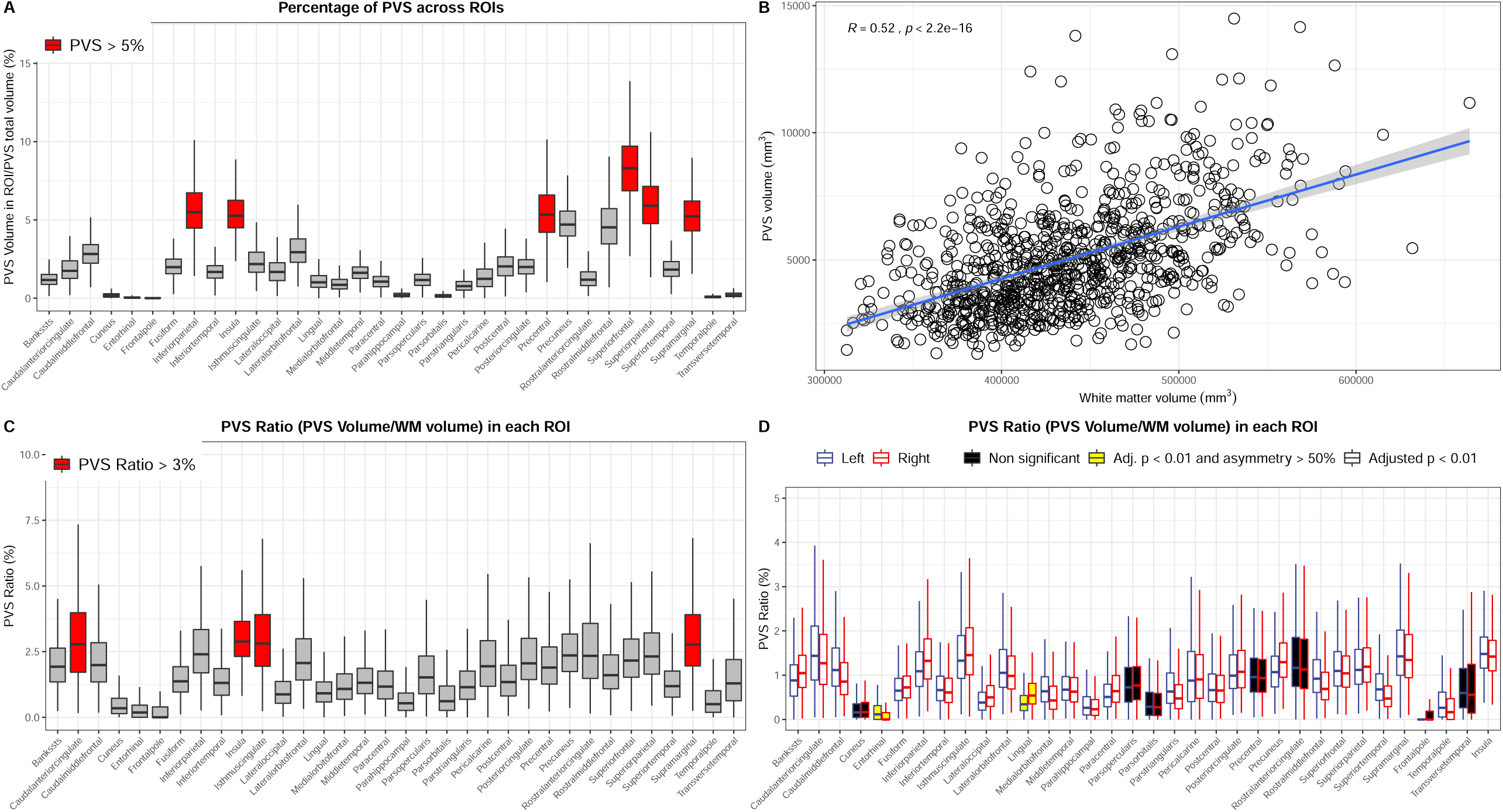
Distribution of the perivascular spaces (PVS) in the white matter and relationship between PVS and white matter. **A.** Boxplot showing the percentage of perivascular space (PVS) in each region of interest (ROI). The ROIs showing more than 5% of total PVS volume are highlighted in red. X-axis labels are white matter regions, parcellated based on Desikan-Killiany atlas using *FreeSurfer* software. “Bankssts”: Banks of the Superior Temporal Sulcus. **B.** Scatterplot showing the significant positive relationship between the measured perivascular space (PVS) and white matter volumes. Spearman’s rank correlation coefficient. **C.** Boxplot showing the PVS ratio (i.e., PVS volume/white matter volume) in each bilateral region of interest (ROI). The reported value in each ROI is the PVS ratio measured on the right and left side of the specific ROI combined. The ROIs with a PVS ratio higher than 3% are highlighted in red. **D.** Boxplot showing the PVS ratio in each unilateral region of interest (ROI). For each ROI, the left boxplot represents the corresponding ROI on the left hemisphere (blue line), while the right boxplot is the corresponding ROI on the right hemisphere (red line). The adjusted p-values refer to the Wilcoxon matched-pairs signed rank test performed in each ROI to compare the two sides. The ROIs with a significant asymmetric distribution of PVS are in white boxes; the ROIs with a significantly asymmetric distribution of PVS having 50% higher PVS ratio on one side compared with its contralateral part are in yellow boxes; the ROIs with a symmetric distribution of PVS ratio across the two hemispheres (i.e., adjusted p-value > 0.01) are in black boxes. Outliers in boxplots are not shown (panels A, B, and D).

We observed a significant relationship between the PVS volume and the measured white matter volume, as assumed *a priori* (*r*=0.52, *p* < 0.0001) (Figure 2B). Therefore, we calculated the PVS ratio, corresponding to the ratio between the PVS volume and the white matter volume.

The average PVS ratio in the whole white matter was 1.14±0.43% (range: 0.34-3.13%). The regions with the highest PVS over white matter volume ratios were the white matter areas adjacent to the cingulate cortex, insula, and supramarginal gyrus, showing a PVS ratio above 3%; on the other hand, the regions with the smallest PVS ratios were the white matter areas underlying the cuneus, entorhinal cortex, and the frontal pole cortex (Figure 2C and supplementary table 2).

When comparing one side of each ROI with its contralateral part in the same subject, the relative difference in PVS ratio was 18% on average, variably exhibiting more PVS on the right or on the left side (Figure 2D). All the regions showed a significant asymmetric distribution of PVS (Wilcoxon matched-pairs, *p*<0.01), except the white matter areas underlying the frontal pole, pars orbitalis and opercularis, anterior cingulate, precentral, transverse temporal, cuneus, and pericalcarine regions (Figure 2D and supplementary table 2). The white matter regions showing on average the highest asymmetric distribution of PVS were those underlying the lingual gyrus (50% higher PVS ratio on the right side) and the entorhinal cortex (120% higher PVS ratio on the left hemisphere) (Figure 2D and supplementary table 2).

Together, these results show that an asymmetric distribution of PVS across the two cerebral hemispheres can be considered physiological in most of the ROIs of healthy adults. Additionally, the entity of the asymmetry can be of great extent in some ROIs, with one side having a PVS ratio up to 120% higher than the contralateral side.

### The PVS ratio is influenced by body mass index, age, and gender

Next, we investigated which demographic and clinical factors affect the amount of PVS in the brain under physiological conditions. A total of 897 participants (503 females and 394 males) were included in the analysis. The mean age was 29.5 in females and 28 in males. The mean body mass index (BMI) was 26.7 kg/m^2^ and was slightly higher in males (26.95) compared with females (26.42). The univariate general linear models testing the clinical factors potentially related with the PVS ratio revealed 4 statistically significant variables: age, BMI, gender and systolic blood pressure (*p*<0.01 in all cases; table 2 and figure 3A-C). Diastolic blood pressure, thyroid stimulating hormone level, hematocrit, and glycated hemoglobin were not significant (Table 2). We included the significant factors as independent variables in a multivariate model testing PVS ratio as dependent variable: higher BMI, older age, and male gender, but not systolic blood pressure, are significant predictors of higher amount of PVS (Table 2). To further analyze the effects of gender and BMI on PVS, we used a two-way ANCOVA model with 4 BMI groups (<20, 20-25, 25-30, >30), adjusting for age. The two-way interaction term between gender and BMI did not reach the statistical significance after controlling for the false discovery rate (supplementary table 3, *p*=0.045). However, on the main effect analyses, we noted that the difference in PVS ratio between males and females is statistically significant in participants with BMI higher than 20 (figure 3D). Interestingly, while in males the relationship between the increase in PVS ratio and the increase in BMI follows a linear trend, in females the increase in PVS is noted exclusively when the BMI is higher than 30 (obese people) (figure 3D). This result suggests that the relationship between BMI and PVS is distinct in males and females and not solely determined by a difference in BMI in the two groups.

**Table 2.** Univariate and multivariate general linear models results. Significant p-values after controlling for the false discovery rate are marked with *.

| Series of univariate general linear models with PVS ratio as dependent variable |  |  |  |
| --- | --- | --- | --- |
|  | Estimate | Confidence Interval (95%) | p-value |
| Age | 0.0162 | 0.008-0.023 | 7.18E-05* |
| BMI | 0.0218 | 0.017-0.027 | 1.46E-15* |
| Gender (male) | 0.1571 | 0.102-0.213 | 1.35E-07* |
| Diastolic blood pressure | 0.0026 | -0.00008-0.0053 | 0.0914 |
| Systolic blood pressure | 0.0039 | 0.002-0.006 | 0.0002* |
| Hematocrit | 0.0031 | -0.003-0.009 | 0.4204 |
| Thyroid Hormone | -0.0012 | -0.031-0.028 | 0.9348 |
| HbA1C | 0.0388 | -0.046-0.123 | 0.4204 |
| Multivariate general linear model with PVS ratio as dependent variable |  |  |  |
|  | Estimate | Confidence Interval (95%) | p-value |
| Age | 0.0206 | 0.010-0.025 | 7.18E-05* |
| BMI | 0.0176 | 0.015-0.026 | 1.46E-15* |
| Gender (male) | 0.1787 | 0.121-0.236 | 1.35E-07* |
| Systolic blood pressure | -0.0009 | -0.003-0.001 | 0.0914 |
| Multivariate general linear model with age-adjusted "Cognitive Function Composite Score" as dependent variable |  |  |  |
|  | Estimate | Confidence Interval (95%) | p-value |
| Gender (male) | 5.6255 | 3.200-8.051 | 6.06E-06* |
| Education | 5.0297 | 4.369-5.691 | < 2E-16* |
| BMI | -0.4131 | -0.648-0.178 | 0.000597* |
| PVS Ratio | -1.7726 | -4.703-1.158 | 0.235489 |
| Multivariate general linear model with age-adjusted "Early Childhood Cognitive Score" as dependent variable |  |  |  |
|  | Estimate | Confidence Interval (95%) | p-value |
| Gender (male) | 2.41763 | 0.404-4.431 | 0.01864* |
| Education | 3.12245 | 2.573-3.672 | < 2E-16* |
| BMI | -0.29904 | -0.495-0.104 | 0.00277* |
| PVS Ratio | -1.95181 | -4.382-0.478 | 0.11527 |

**Figure 3.**
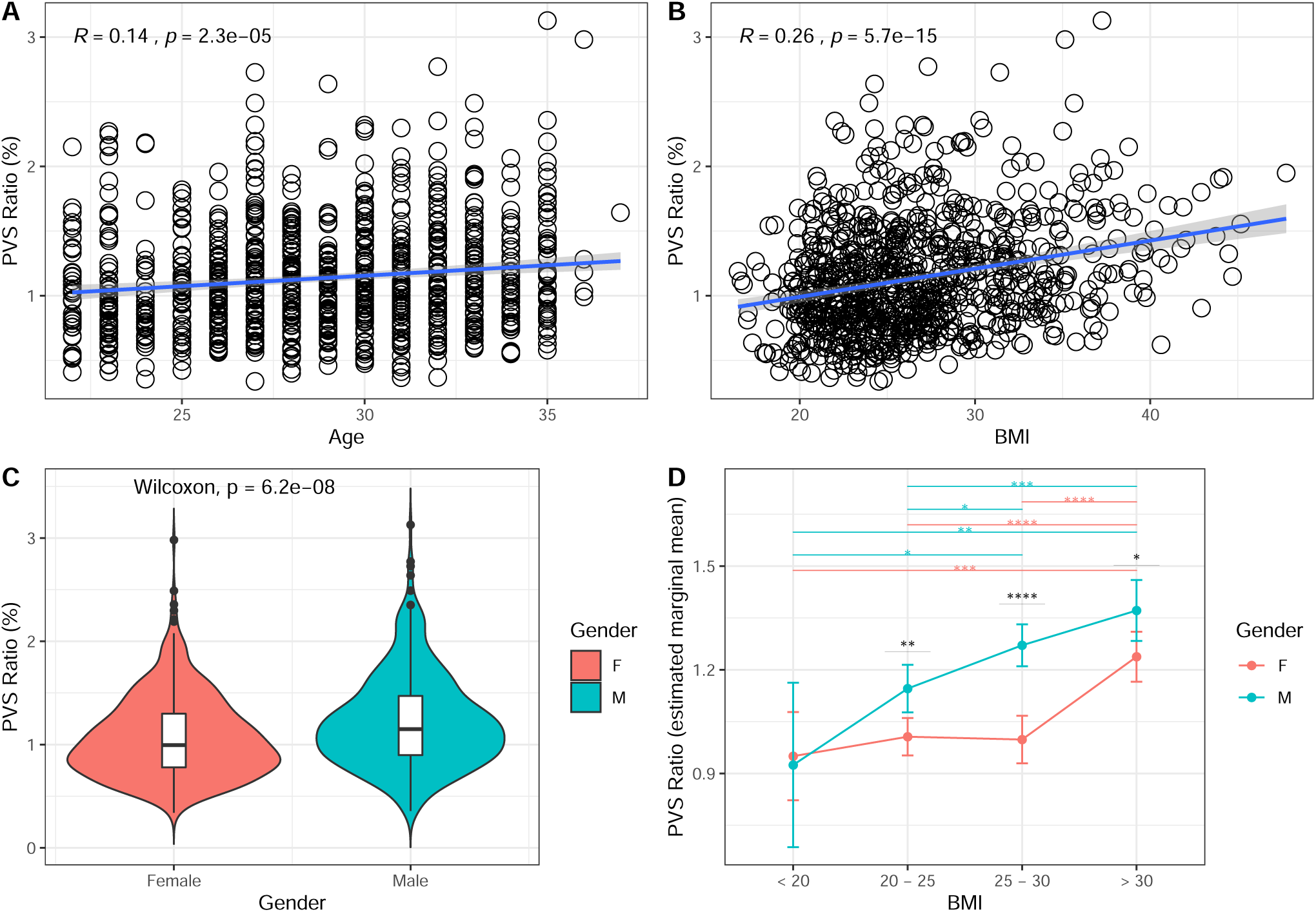
The perivascular space (PVS) ratio is influenced by age, body mass index (BMI), and gender. Scatterplots showing the relationship between PVS and age (**A**) and BMI (**B**) (Spearman’s rank correlation coefficient). **C.** Violin plot showing the statistically significant difference in PVS ratio between males (green) and female (red) participants (Wilcoxon rank sum test). **D.** Estimated marginal means of PVS ratio in males and females represented in each BMI group. Significance by ANCOVA for main effects (black *) and post-hoc comparisons (green and red *) controlling for age. The error bars are lower and upper bounds on a 95% confidence interval of the estimate. *: adjusted p-value < 0.05; **: adjusted p-value < 0.01; ***: adjusted p-value < 1×10^−3^; ****: adjusted p-value < 1×10^−4^. The following post-hoc comparisons were not significant after controlling for the false discovery rate: PVS ratio difference in males between BMI groups “< 20” and “20-25”; PVS ratio difference in males between BMI groups “< 20” and “25-30”.

We also investigated the role that cigarette smoking and alcohol could play in modulating PVS. The PVS ratio was not significantly different in regular smokers, occasional smokers, and non-smokers (Kruskal-Wallis, *p*=0.49), and the number of cigarettes per day did not significantly correlate with the PVS ratio (*r*=-0.04, *p*=0.55). The total number of alcoholic drinks consumed in one week on average was not significantly correlated with the PVS ratio (*r*=0.55, *p*=0.1). In summary, this analysis shows that age, gender, and BMI influence the total volume of PVS, and that the relationship between BMI and PVS is different in males versus females.

### Cognitive functions in healthy adults are not influenced by PVS

Whether the occurrence of enlarged PVS on MRI in the general elderly population is associated with cognitive dysfunction remains unclear(56,57). Here we analyzed the effect of the PVS ratio to cognition in healthy young adults. The average years of education in this population are 14.9±1.8 (range: 11-17) and the mean Mini-Mental State Examination (MMSE) is 29±1 (range: 23-30); the education level is slightly higher in females (14.98) compared with males (14.77), and MMSE is not significantly different in females compared with males (29.05 and 28.96, respectively; Wilcoxon, *p*=0.44). The PVS ratio is not significantly correlated with the level of education (*r*=-0.04, *p*=0.24) and the MMSE (*r*=0.01, *p*=0.73).

We performed a principal component analysis on a set of 19 NIH Toolbox age-adjusted behavioral tests to identify the tests explaining most of the variance: within the first component, explaining 30% of the variance, the most influential tests (loadings > 0.35) are the Cognitive Function Composite score (loading: 0.40) and the Early Childhood Composite score (loading: 0.37). These age-adjusted scores were included in a linear model (each at a time), corrected by gender and education, as dependent variables to investigate whether the PVS ratio affects cognitive performance. The models showed a significant trend towards the PVS ratio as a factor affecting both the Cognitive Function Composite score and the Early Childhood Composite score (*p*=0.0321 and *p*=0.0150, respectively). However, when BMI was added as a covariate in both models, PVS ratio did not reach the statistical significance, while the BMI was found to be a significant factor for both the analyzed cognitive scores (*p*<0.01 in both cases), where a higher BMI was associated with lower scores (Table 2). These results suggest that a higher amount of PVS in the brain of young adults does not significantly affect cognition, and that higher BMI is associated with lower cognitive scores. Therefore, the apparent association between the greater amount of PVS and worse cognitive performance in a healthy young population is potentially caused by the linear relationship between BMI and PVS.

### The PVS ratio is influenced by the time of day

Next, we investigated whether the sleep quality and quantity as well as the time of day play a role in the extent of PVS detectable on MRI. In the whole cohort (n=897), we did not find a significant relationship between PVS ratio and the average number of hours of sleep (*r*=-0.05, *p*=0.11) or the sleep quality index (*r*=0.04, *p*=0.2). Since in our population we found a high level of inter-subject variability in PVS ratio, we selectively analyzed 45 participants (31 females, 14 males, mean age: 30.3±3.3) that underwent a second MRI scan, with the same scanner and protocol, after 139±69 days. The mean BMI (26.9±5.8) and amount of sleep (7.1±0.9 hours) before the first MRI scan were not significantly different from those before the second MRI session (26.6±5.7 and 7.2±0.9, *p*=0.23 and 0.41, respectively). The intra-individual difference in PVS volume and the corresponding difference in minutes between the MRI scan performed at a later time of day and the MRI scan performed at an earlier time of day was computed (figure 4A). We found a statistically significant relationship between the time difference and the PVS volume change (*r*=0.34, *p*=0.022. Figure 4B): the increase in PVS volume was greater when the difference between the time-of-day of the two MRI scans was larger. These results suggest that, in people with stable sleep habits, the amount of fluid within the PVS physiologically changes throughout the day, with more fluid detectable at later times of the day.

**Figure 4.**
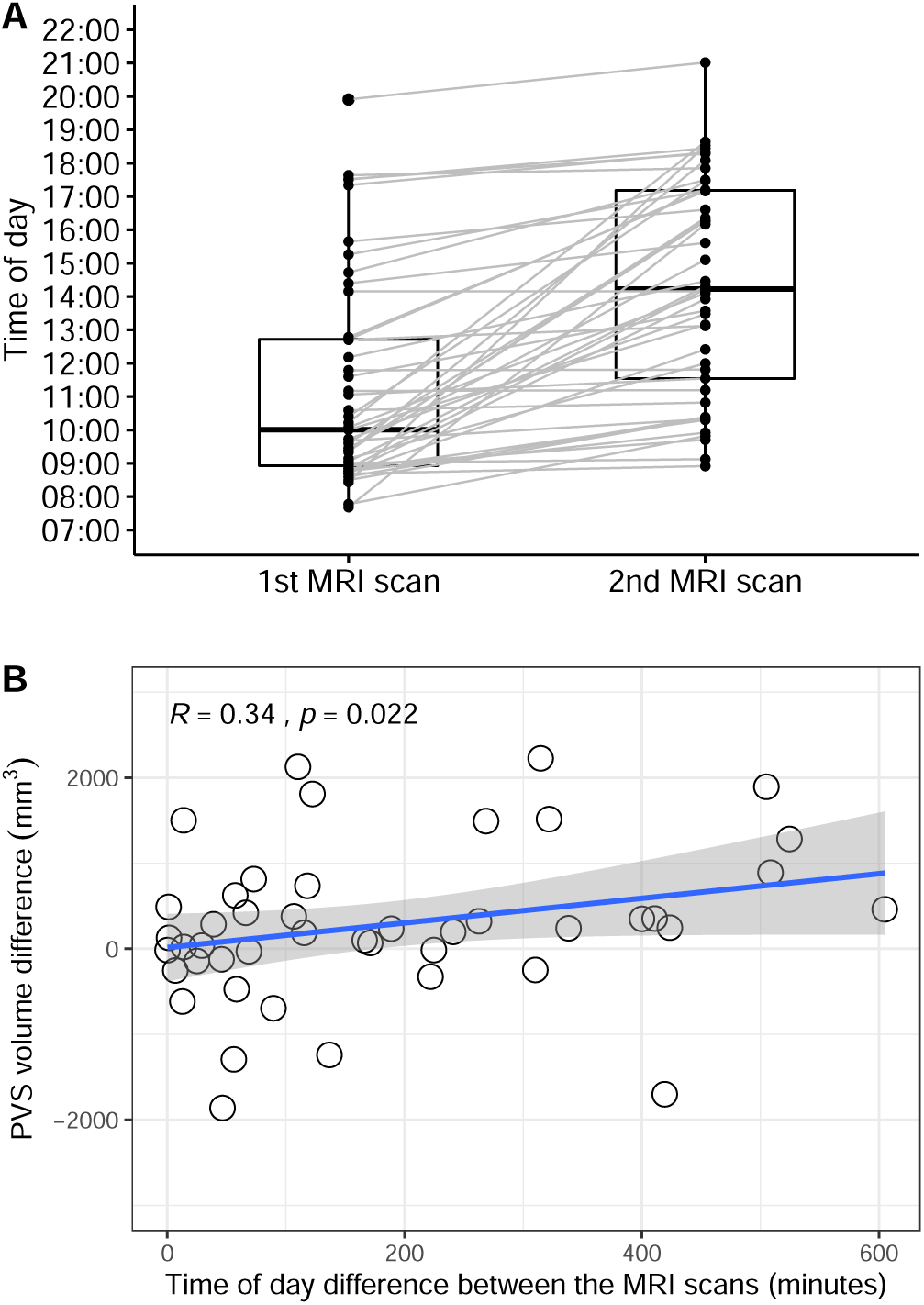
The perivascular space (PVS) volume in the single individual changes throughout the day. **A.** Boxplot showing the difference in time of day between the first MRI scan and the second MRI scan in each participant (n=45). **B.** Scatterplot showing the relationship between the difference in time-of-day the two MRI scans have been performed (in minutes) and the corresponding changes measured in the perivascular space volume (PVS). Spearman’s rank correlation coefficient. None of the values included in this plot is a significant outlier (Extreme studentized deviate method, *p*>0.01).

### The PVS ratio is influenced by genetic factors

Finally, to study the relationship between PVS and genetic factors, we focused on 3 groups: 51 couples of monozygotic twins (62 females and 40 males, mean age: 29.3±3.4), 29 couples of dizygotic twins (36 females and 22 males, mean age: 29.3±3.3), and 143 couples of non-twin siblings (148 females and 138 males, mean age: 28.4±3.9) available on the Human Connectome Project dataset. The correlation between the PVS ratio of each participant with the PVS ratio of the corresponding sibling was statistically significant in monozygotic twins and non-twin siblings (*p*<0.01 in both cases, figure 5A and 5C), but did not reach statistical significance in dizygotic twins after controlling for the false discovery rate, possibly due to the lower sample size (*p*=0.037. Figure 5B). The correlation was still significant when all couples of siblings (twins and non-twins) were grouped together (*r*=0.54, *p*<0.01). After randomization of the pairs, achieved by exchanging one member of the siblings with another member from a different couple, the correlation between the 2 PVS ratios in the new randomized couples was not significant anymore (*r*=0.027, *p*=0.64, figure 5D). In any of the 3 groups, the difference in the PVS ratio measured across matched siblings was not correlated with the corresponding difference in BMI between each member of the pairs (*p*=0.91, 0.34, and 0.97, in monozygotic, dizygotic, and non-twin siblings, respectively. Supplementary figure 1). These results suggest that genetic factors influence the amount of PVS in the brain.

**Figure 5.**
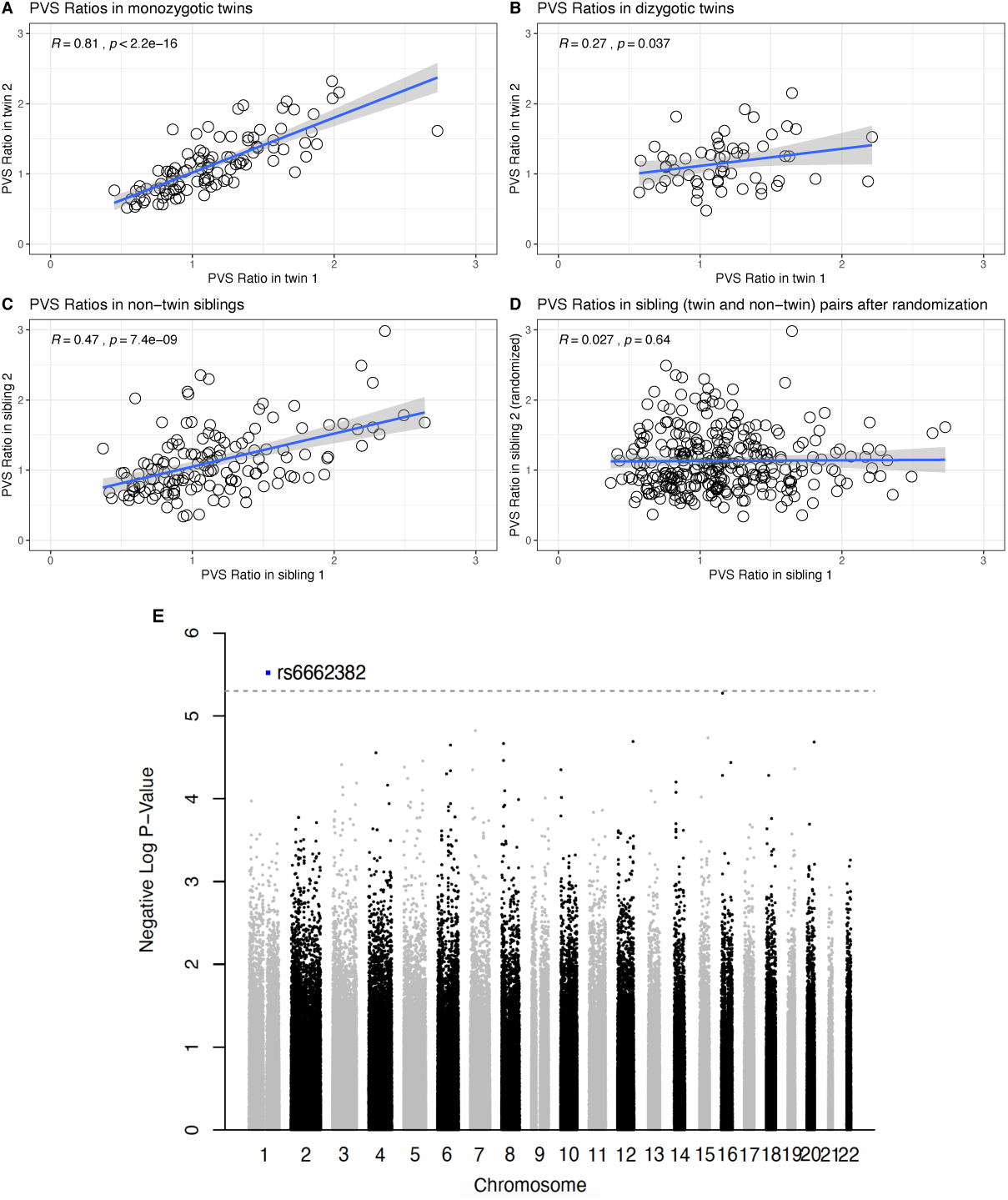
The perivascular space (PVS) ratio is influenced by genetics. Scatterplots showing the relationship of the PVS ratio in each member of the couples plotted against the PVS ratio of the corresponding sibling, in monozygotic twins (**A**), dizygotic twins (**B**), and non-twin siblings (**C**). **D**. The correlation is not significant after randomization of one member in each couple, including twins and non-twin siblings. Spearman’s rank correlation coefficient. **E**. Manhattan plot showing the association *p*-values between SNPs and PVS ratio across the genome.

To gain insights on the specific genetic elements that could affect PVS, we performed a genome-wide association analysis in the 831 participants for which genetic data was available, with the goal of finding SNPs associated with PVS ratio. A SNP located in the OR10T2 gene (Olfactory Receptor Family 10 Subfamily T Member 2) in chromosome 1 was found to be significantly associated with PVS ratio at a suggestive association threshold (p=3E-6. Figure 5E).

## Discussion

Our findings demonstrate that perivascular spaces display a significant inter-subject variability in a healthy young population (range: 1.31-14.49 cm^3^) and that several factors contribute to the amount of PVS measured on MRI. We confirmed that the absolute volume of PVS is strongly correlated with the white matter volume, corroborating the importance of computing a white-matter-adjusted measure of PVS (PVS ratio). This is usually impracticable in the analysis performed with visual rating scales, representing a significant limiting factor for the correct interpretation of the results they can provide.

Concerning the distribution of PVS in the brain, the centrum semiovale and basal ganglia are typically recognized as the area where most of the PVS are usually visible(58), but the physiological regional division in the white matter is not known. We showed that the majority of PVS are visible in the white matter below the superior frontal and parietal cortices, while the highest PVS ratio was found in the white matter adjacent to the cingulate and insular cortices (*capsulae extrema* and *externa*). Moreover, an asymmetric distribution of PVS across the two hemispheres was found in our healthy population, with some regions presenting more than 50% times higher PVS ratio on one side compared with the contralateral part. This is particularly relevant for stroke and post-traumatic epilepsy research, since the asymmetry in perivascular flow seems to play a key pathogenetic role in those diseases(19,59). To our knowledge, this neuroimaging-based PVS map is the most structurally complete atlas of the human PVS to date and can be used as a reference for future quantitative investigations of PVS. We also investigated factors potentially affecting PVS volume in a healthy population.

While aging has already been shown to be associated with enlarged PVS(60), we confirmed this finding even in a population with a relatively narrow age range (22-37). Interestingly, BMI represents a novel factor influencing the amount of PVS: BMI was the most significant variable correlated with PVS ratio in our population. Previous studies have shown that BMI has a linear relationship with CSF pressure in a population with normal CSF pressure values (8-15 mmHg)(61). Even though it was not possible to measure the CSF pressure in our population, this result suggests that the correlation between PVS and BMI could be a consequence of higher CSF pressure in participants with high BMI.

Additionally, obesity is known to critically affect vascular function, including the vascular contractile response(62), which is thought to be one of the main factors driving fluid movement through the PVS(63). Hence, vascular contractility could represent another link between BMI and PVS. Nevertheless, since BMI is a non-specific index which can be equally influenced by lean body mass, fat, and body fluid, the biological mechanisms explaining the relationship between BMI and PVS remain to be investigated.

Another interesting finding is that males showed higher PVS ratio than females. Previous studies using visual rating scores have reported greater prevalence of enlarged PVS in men compared with women, both in normal elderly and dementia cohorts(10,31,64). Intriguingly, we observed an age-corrected gender difference in PVS ratio in all BMI groups except in participants with BMI less than 20 and the effect of BMI on PVS ratio is more pronounced in males compared with females, especially in people with BMI between 20 and 30. BMI is positively correlated with plasma biomarkers of inflammation(65). The astrocytic response to inflammation has been previously demonstrated to be higher in males compared with females, possibly due to the perinatal testosterone which programs astrocytes for a different response to inflammatory challenges(66). Therefore, the higher amount of PVS we found in male participants compared with BMI-matched females when BMI is higher than 20 might be related to a more vigorous inflammatory response in males, which can affect the perivascular flow and the size of PVS(66). It would be interesting for future studies to explore the effects of high-fat diet on the cerebrovasculature and the perivascular flow, comparing males and females, and to verify the potentially different changes in the perivascular flow and whether these changes lead to pathological modifications at the cellular and cognitive levels.

Regarding cognitive function, our results do not substantiate a significant relationship between neuropsychological test scores and PVS ratio in young adults, although there is a trend showing more PVS in people who scored worse in some cognitive tests. Remarkably, this trend is mostly explained by the BMI, which appeared as a critical factor in relation with some cognitive scores, where a higher BMI was associated with lower scores. This finding, however, cannot support a biological causal connection between BMI and specific aspects of cognition. For example, it should be noted that in our population higher BMI was inversely correlated with the level of education (supplementary figure 2). In fact, several other epidemiological and cultural factors, potentially associated with BMI and not included in our analysis, could at least partially affect the scores obtained in the neuropsychological tests administered.

Concerning the analysis of sleep, previous human studies showed that people with impaired sleep efficiency and obstructive sleep apnea present increased PVS visibility (measured with visual rating scales), which was indirectly interpreted as PVS dysfunction(15,67). Our findings do not show any significant difference in PVS ratio between people on different hours of sleep. This might be due not only to the high inter-subject variability on PVS related to other factors, but also to different body postures exhibited by each participant during sleep(68), which were not considered in our study. In fact, animal studies showed that the position assumed during sleep is another critical factor affecting the CSF transport in the PVS(68). Additionally, influence of sleep problems on the brain has been reported to occur in midlife and older ages(69,70), so the effects on PVS might still be undetectable in the young population that we analyzed. On the other hand, our results showed the time-of-day as an important element affecting the PVS volume. Specifically, a subset of people that underwent MRI scans twice in different days at different time, showed a significantly higher PVS volume in the afternoon and the evening compared with the PVS volume measured on the same individual’ scan acquired at an earlier time of day, which is indicative of a circadian fluctuation in the perivascular flow. Increased uptake of CSF tracer gadobutrol into the entorhinal cortex overnight has been recently shown in patients with idiopathic normal pressure hydrocephalus and controls(71,72), suggesting a critical role of natural sleep for glymphatic function, as indicated by studies in rodents(73). A similar fluctuation has been also demonstrated in diffusivity measures of brain tissue derived from diffusion tensor imaging: mean diffusivity, which is significantly influenced by perivascular spaces as well(74), was found to systematically increase from morning to afternoon scans(75). Here we further validated these findings, showing an increased amount of fluid within the PVS in the white matter at later time of day in the same person. It is possible that these changes are related to circadian oscillations in blood pressure and/or respiration, two regulators of the perivascular flow(63,76).

Finally, we analyzed for the first time the influence of genetic factors on the PVS ratio in a healthy population. We found that couples of siblings have more similar PVS ratio compared with couples of non-siblings. The similarity was more pronounced in monozygotic twins and was not explained by the difference in BMI. Subsequently, we looked for genome-wide significant association between SNPs and PVS ratio. Although none of the SNPs passed the Bonferroni threshold for GWAS, possibly because of the relatively small sample size of the analyzed cohort, SNPs that are associated with suggestive significance also provide crucial biological insights, given the polygenic and multifactorial nature of many complex phenotypes such as PVS(77). Interestingly, the SNP showing the most significant association with PVS ratio was located in the OR10T2 gene, a highly conserved region which encodes for one type of olfactory receptors. Previous studies in humans and mammalians have shown an intimate connection between CSF circulation and olfactory-associated perineural, perivascular, and lymphatic compartments, which represent a significant drainage pathway and access route to the brain(78–82). Olfactory receptors may therefore represent an important regulator of the inflow and outflow of molecules in the perivascular spaces, consequently affecting the amount of fluid within the PVS.

Of note, since the measure and detectability of perivascular spaces depend on the resolution of the images and the field strength of the MRI system(6), a direct comparison of our results with data acquired at different field strength and/or resolution might be inappropriate. This study represents the largest quantitative analysis of PVS in humans using MRI and the only one performed in healthy young adults. These findings can be used as a reference atlas for clinicians and researchers investigating PVS: we provide PVS volumes, PVS/white matter ratios, and their regional distribution that can be helpful when studying PVS under pathological conditions and when attempting to identify patients with abnormal PVS size, location, and asymmetry. Moreover, we report several novel factors that significantly contribute to the observed high inter-subject variability of PVS visibility in healthy participants, that should be taken into consideration in future research studies analyzing PVS.

## Acknowledgements

Data were provided in part by the Human Connectome Project, WU-Minn Consortium (Principal Investigators: David Van Essen and Kamil Ugurbil; 1U54MH091657) funded by the 16 NIH Institutes and Centers that support the NIH Blueprint for Neuroscience Research; and by the McDonnell Center for Systems Neuroscience at Washington University. MRI scans and clinical data from the Human Connectome Project can be accessed from https://www.humanconnectome.org. F.S. was partly supported by R01NS100973. The authors would like to thank Lucia Ichino for her critical evaluation of the manuscript.

## Author contribution statement

G.B., F.S., N.S.B, M.L. and A.W.T. designed the research study. G.B., F.S., and N.S.B. analyzed and interpreted the data. F.S., N.S.B, M.L. and A.W.T. provided critical reading of the manuscript and G.B. wrote the manuscript. All authors edited and revised the manuscript and approved final submission.

---

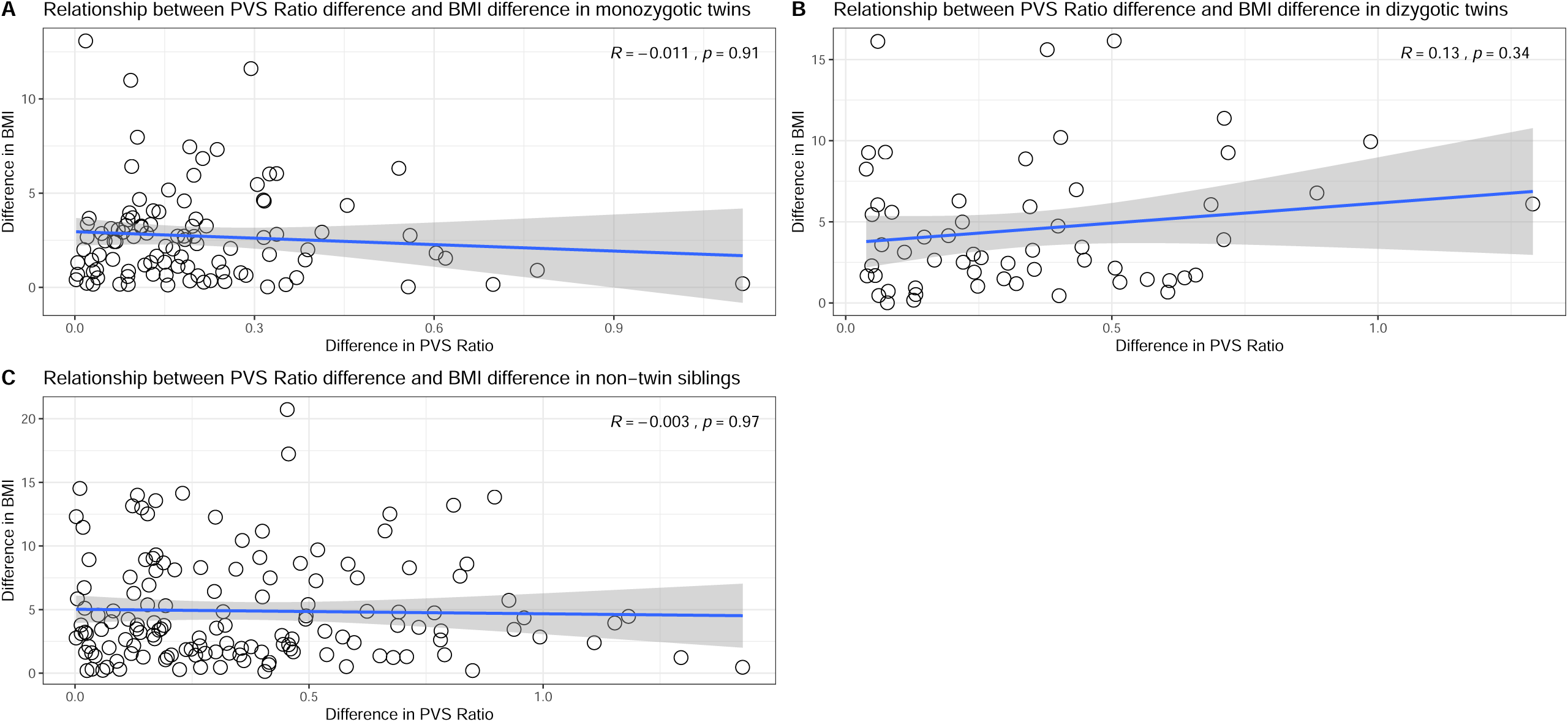

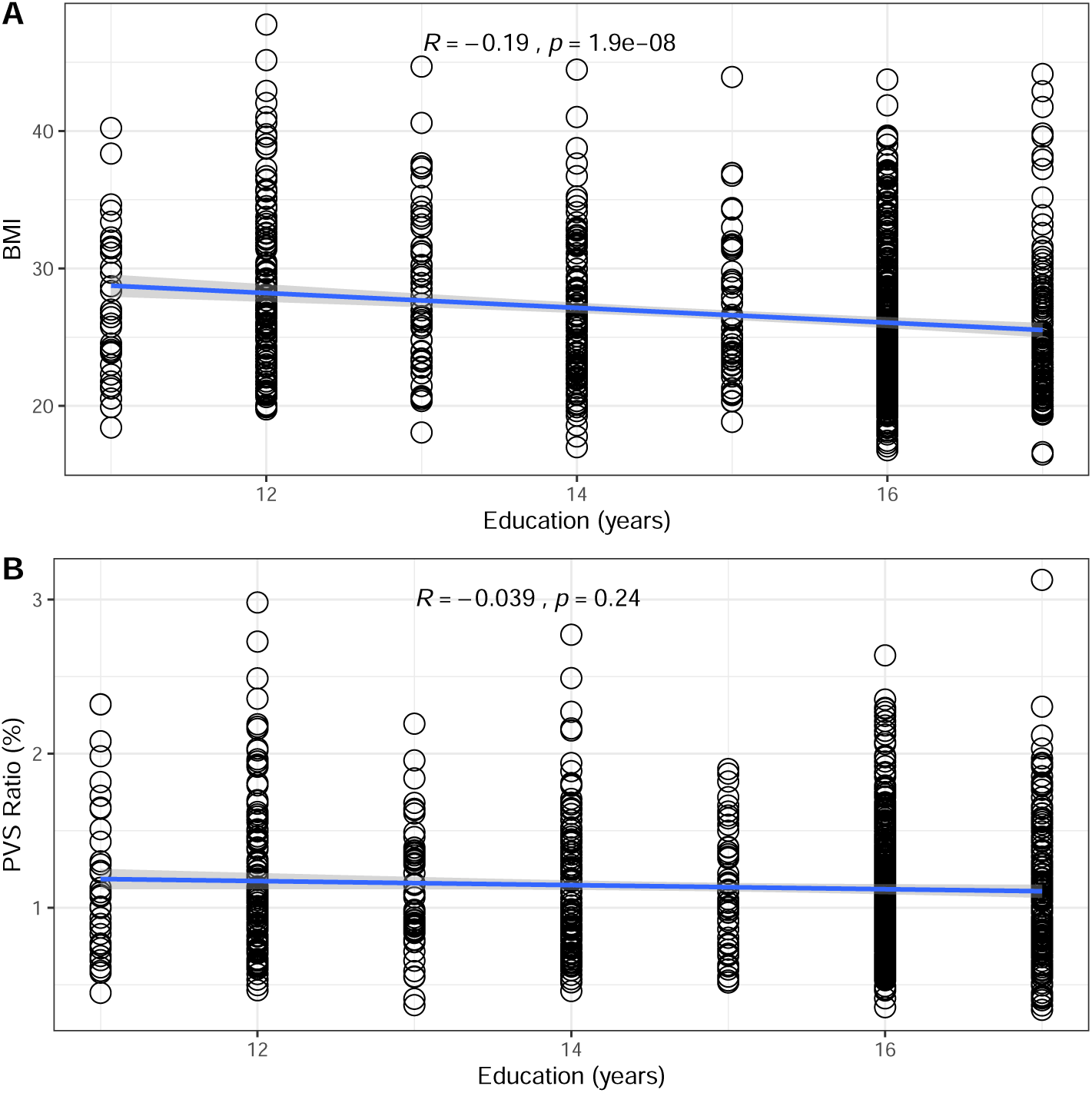

## References

1. Zhang ET, Inman CB, Weller RO. Interrelationships of the pia mater and the perivascular (Virchow-Robin) spaces in the human cerebrum. J Anat [Internet]. 1990;170:111–23. Available from: http://www.pubmedcentral.nih.gov/articlerender.fcgi?artid=1257067&tool=pmcentrez&rendertype=abstract

2. Rasmussen MK, Mestre H, Nedergaard M. The glymphatic pathway in neurological disorders. Lancet Neurol [Internet]. 2018 Nov 1 [cited 2019 May 27];17(11):1016–24. Available from: https://www-sciencedirect-com.libproxy2.usc.edu/science/article/pii/S1474442218303181?via%3Dihub

3. Tarasoff-Conway JM, Carare RO, Osorio RS, Glodzik L, Butler T, Fieremans E, et al. Clearance systems in the brain—implications for Alzheimer disease. Nat Rev Neurol. 2015;11(8):457.

4. Wardlaw JM, Valdés Hernández MC, Muñoz-Maniega S. What are white matter hyperintensities made of? Relevance to vascular cognitive impairment. J Am Heart Assoc. 2015;4(6):e001140.

5. Brown R, Benveniste H, Black SE, Charpak S, Dichgans M, Joutel A, et al. Understanding the role of the perivascular space in cerebral small vessel disease. Cardiovasc Res [Internet]. 2018 Sep 1 [cited 2019 Aug 21];114(July):1–9. Available from: https://academic.oup.com/cardiovascres/article/114/11/1462/4991897

6. Barisano G, Sepehrband F, Ma S, Jann K, Cabeen R, Wang DJ, et al. Clinical 7T MRI: are we there yet? A review about magnetic resonance imaging at ultra-high field. Br J Radiol. 2018 Oct;91:20180492.

7. Wardlaw JM, Benveniste H, Nedergaard M, Zlokovic Ef B V, Mestre H, Lee H, et al. Perivascular Spaces in the Brain: Anatomy, Physiology, and Contributions to Pathology of Brain Diseases. Vol. 16, Nature Reviews Neurology. Nature Research; 2020 Mar.

8. Cai K, Tain R, Das S, Damen FC, Sui Y, Valyi-Nagy T, et al. The feasibility of quantitative MRI of perivascular spaces at 7T. J Neurosci Methods [Internet]. 2015;256:151–6. Available from: http://dx.doi.org/10.1016/j.jneumeth.2015.09.001

9. Zong X, Park SH, Shen D, Lin W. Visualization of perivascular spaces in the human brain at 7T: Sequence optimization and morphology characterization. Neuroimage [Internet]. 2016;125:895–902. Available from: http://dx.doi.org/10.1016/j.neuroimage.2015.10.078

10. Zhu YC, Dufouil C, Mazoyer B, Soumaré A, Ricolfi F, Tzourio C, et al. Frequency and location of dilated Virchow-Robin spaces in elderly people: A population-based 3D MR imaging study. Am J Neuroradiol. 2011 Apr;32(4):709–13.

11. Maclullich AMJ, Wardlaw JM, Ferguson KJ, Starr JM, Seckl JR, Deary IJ. Enlarged perivascular spaces are associated with cognitive function in healthy elderly men. J Neurol Neurosurg Psychiatry [Internet]. 2004 Nov 1 [cited 2017 Sep 25];75(11):1519–23. Available from: http://www.ncbi.nlm.nih.gov/pubmed/15489380

12. Taber KH, Shaw JB, Loveland KA, Pearson DA, Lane DM, Hayman LA. Accentuated Virchow-Robin Spaces in the Centrum Semiovale in Children with Autistic Disorder. J Comput Assist Tomogr. 2004;28(2):263–8.

13. Rollins NK, Deline C, Morriss MC. Prevalence and Clinical Significance of Dilated Virchow-Robin Spaces in Childhood. Radiology. 1993;189:53–7.

14. Patankar TF, Baldwin R, Mitra D, Jeffries S, Sutcliffe C, Burns A, et al. Virchow-Robin space dilatation may predict resistance to antidepressant monotherapy in elderly patients with depression. J Affect Disord. 2007;97(1–3):265–70.

15. Berezuk C, Ramirez J, Gao F, Scott CJM, Huroy M, Swartz RH, et al. Virchow-Robin Spaces: Correlations with Polysomnography-Derived Sleep Parameters. Sleep [Internet]. 2015 Jun 1 [cited 2017 Sep 25];38(6):853–8. Available from: https://academic.oup.com/sleep/article-lookup/doi/10.5665/sleep.4726

16. Achiron A, Faibel M. Sandlike appearance of Virchow-Robin spaces in early multiple sclerosis: A novel neuroradiologic marker. Am J Neuroradiol. 2002;23(3):376–80.

17. Wuerfel J, Haertle M, Waiczies H, Tysiak E, Bechmann I, Wernecke KD, et al. Perivascular spaces--MRI marker of inflammatory activity in the brain? Brain [Internet]. 2008 Aug 21 [cited 2017 Sep 25];131(9):2332–40. Available from: https://academic.oup.com/brain/article-lookup/doi/10.1093/brain/awn171

18. Inglese M, Bomsztyk E, Gonen O, Mannon LJ, Grossman RI, Rusinek H. Dilated Perivascular Spaces: Hallmarks of Mild Traumatic Brain Injury. Am J Neuroradiol. 2005;26(4).

19. Duncan D, Barisano G, Cabeen R, Sepehrband F, Garner R, Braimah A, et al. Analytic Tools for Post-traumatic Epileptogenesis Biomarker Search in Multimodal Dataset of an Animal Model and Human Patients. Front Neuroinform [Internet]. 2018;12:86. Available from: https://www.frontiersin.org/article/10.3389/fninf.2018.00086

20. Laitinen L V., Chudy D, Tengvar M, Hariz MI, Tommy Bergenheim A. Dilated perivascular spaces in the putamen and pallidum in patients with Parkinson’s disease scheduled for pallidotomy: A comparison between MRI findings and clinical symptoms and signs. Mov Disord. 2000;15(6):1139–44.

21. Di Costanzo A, Di Salle F, Santoro L, Bonavita V, Tedeschi G. Dilated Virchow-Robin spaces in myotonic dystrophy: Frequency, extent and significance. Eur Neurol. 2001;46(3):131–9.

22. Miyata M, Kakeda S, Iwata S, Nakayamada S, Ide S, Watanabe K, et al. Enlarged perivascular spaces are associated with the disease activity in systemic lupus erythematosus. Sci Rep. 2017 Dec 1;7(1):1–10.

23. Potter GM, Doubal FN, Jackson CA, Chappell FM, Sudlow CL, Dennis MS, et al. Enlarged perivascular spaces and cerebral small vessel disease. Int J Stroke [Internet]. 2015 Apr [cited 2017 Sep 21];10(3):376–81. Available from: http://www.ncbi.nlm.nih.gov/pubmed/23692610

24. Rouhl RPW, van Oostenbrugge RJ, Knottnerus ILH, Staals JEA, Lodder J. Virchow-Robin spaces relate to cerebral small vessel disease severity. J Neurol [Internet]. 2008;255(5):692–6. Available from: http://link.springer.com/10.1007/s00415-008-0777-y

25. Ohba H, Pearce L, Potter G, Benavente O. Enlarged perivascular spaces in lacunar stroke patients. The secondary prevention of small subcortical stroked (SPS3) trial. In: International Stroke Conference. 2012. p. A151.

26. Doubal FN, MacLullich AMJ, Ferguson KJ, Dennis MS, Wardlaw JM. Enlarged perivascular spaces on MRI are a feature of cerebral small vessel disease. Stroke [Internet]. 2010 Mar 1 [cited 2017 Sep 25];41(3):450–4. Available from: http://www.ncbi.nlm.nih.gov/pubmed/20056930

27. Wardlaw JM, Smith EE, Biessels GJ, Cordonnier C, Fazekas F, Frayne R, et al. Neuroimaging standards for research into small vessel disease and its contribution to ageing and neurodegeneration. Lancet Neurol [Internet]. 2013 Aug [cited 2017 Dec 26];12(8):822–38. Available from: http://www.ncbi.nlm.nih.gov/pubmed/23867200

28. Charidimou A, Jaunmuktane Z, Baron J-C, Burnell M, Varlet P, Peeters A, et al. White matter perivascular spaces: an MRI marker in pathology-proven cerebral amyloid angiopathy? Neurology [Internet]. 2014 Jan 7 [cited 2017 Sep 25];82(1):57–62. Available from: http://www.ncbi.nlm.nih.gov/pubmed/24285616

29. Martinez-Ramirez S, Pontes-Neto OM, Dumas AP, Auriel E, Halpin A, Quimby M, et al. Topography of dilated perivascular spaces in subjects from a memory clinic cohort. Neurology. 2013;80(17):1551–6.

30. Roher AE, Kuo Y-M, Esh C, Knebel C, Weiss N, Kalback W, et al. Cortical and leptomeningeal cerebrovascular amyloid and white matter pathology in Alzheimer’s disease. Mol Med [Internet]. 2003 [cited 2017 Dec 26];9(3–4):112–22. Available from: http://www.ncbi.nlm.nih.gov/pubmed/12865947

31. Ramirez J, Berezuk C, Mcneely AA, Scott CJM, Gao F, Black SE. Visible Virchow-Robin Spaces on Magnetic Resonance Imaging of Alzheimer’s Disease Patients and Normal Elderly from the Sunnybrook Dementia Study. J Alzheimer’s Dis [Internet]. 2015 [cited 2018 Jan 9];43:415–24. Available from: https://content.iospress.com/download/journal-of-alzheimers-disease/jad132528?id=journal-of-alzheimers-disease%2Fjad132528

32. Woollam DH, Millen JW. The perivascular spaces of the mammalian central nervous system and their relation to the perineuronal and subarachnoid spaces. J Anat [Internet]. 1955 Apr [cited 2020 Feb 5];89(2):193–200. Available from: http://www.ncbi.nlm.nih.gov/pubmed/14367214

33. Van Essen DC, Ugurbil K, Auerbach E, Barch D, Behrens TEJ, Bucholz R, et al. The Human Connectome Project: A data acquisition perspective. Neuroimage. 2012;62(4):2222–31.

34. Van Essen DC, Smith SM, Barch DM, Behrens TEJ, Yacoub E, Ugurbil K. The WU-Minn Human Connectome Project: An overview. Neuroimage. 2013 Oct 15;80:62–79.

35. Hu LS, Baxter LC, Smith KA, Feuerstein BG, Karis JP, Eschbacher JM, et al. Relative Cerebral Blood Volume Values to Differentiate High-Grade Glioma Recurrence from Posttreatment Radiation Effect: Direct Correlation between Image-Guided Tissue Histopathology and Localized Dynamic Susceptibility-Weighted Contrast-Enhanced Perfusio. Am J Neuroradiol. 2008 Dec;30(3):552–8.

36. Folstein MF, Robins LN, Helzer JE. The mini-mental state examination. Arch Gen Psychiatry. 1983;40(7):812.

37. Buysse DJ, Reynolds CF, Monk TH, Berman SR, Kupfer DJ. The Pittsburgh sleep quality index: A new instrument for psychiatric practice and research. Psychiatry Res. 1989 May 1;28(2):193–213.

38. Glasser MF, Sotiropoulos SN, Wilson JA, Coalson TS, Fischl B, Andersson JL, et al. The minimal preprocessing pipelines for the Human Connectome Project. Neuroimage [Internet]. 2013 Oct 15 [cited 2019 Dec 16];80:105–24. Available from: https://linkinghub.elsevier.com/retrieve/pii/S1053811913005053

39. Jenkinson M, Beckmann CF, Behrens TEJ, Woolrich MW, Smith SM. Fsl. Neuroimage. 2012;62(2):782–90.

40. Fischl B. FreeSurfer. Neuroimage [Internet]. 2012 Aug 15 [cited 2019 Dec 16];62(2):774–81. Available from: http://www.ncbi.nlm.nih.gov/pubmed/22248573

41. Sepehrband F, Barisano G, Sheikh-Bahaei N, Cabeen RP, Choupan J, Law M, et al. Image processing approaches to enhance perivascular space visibility and quantification using MRI. Sci Rep [Internet]. 2019 Dec 26 [cited 2019 Aug 26];9(1):12351. Available from: http://www.nature.com/articles/s41598-019-48910-x

42. Manjón J V., Coupé P, Martí-Bonmatí L, Collins DL, Robles M. Adaptive non-local means denoising of MR images with spatially varying noise levels. J Magn Reson Imaging. 2010;31(1):192–203.

43. Avants BB, Tustison N, Song G. Advanced Normalization Tools (ANTS). Insight J. 2009;1–35.

44. Avants BB, Tustison NJ, Wu J, Cook PA, Gee JC. An open source multivariate framework for N-tissue segmentation with evaluation on public data. Neuroinformatics. 2011 Dec;9(4):381–400.

45. Frangi AF, Niessen WJ, Vincken KL, Viergever MA. Multiscale vessel enhancement filtering. Med Image Comput Comput Interv Miccai’98 1496, [Internet]. 1998 Oct 11 [cited 2017 Sep 25];1496:130–7. Available from: http://link.springer.com/10.1007/BFb0056195

46. Cabeen RP, Laidlaw DH, Toga AW. Quantitative Imaging Toolkit: Software for Interactive 3D Visualization, Data Exploration, and Computational Analysis of Neuroimaging Datasets. In: Proceedings of the Joint Annual Meeting ISMRM-ESMRMB. Paris, France; 2018. p. 8882.

47. Hou Y, Park SH, Wang Q, Zhang J, Zong X, Lin W, et al. Enhancement of Perivascular Spaces in 7 T MR Image using Haar Transform of Non-local Cubes and Block-matching Filtering. Sci Rep [Internet]. 2017;7(1):1–12. Available from: http://dx.doi.org/10.1038/s41598-017-09336-5

48. Zhang J, Gao Y, Park SH, Zong X, Lin W, Shen D. Structured Learning for 3-D Perivascular Space. 2017;64(12):2803–12.

49. Ballerini L, Lovreglio R, Valdés Hernández MDC, Ramirez J, MacIntosh BJ, Black SE, et al. Perivascular Spaces Segmentation in Brain MRI Using Optimal 3D Filtering. Sci Rep. 2018;8(1):1–11.

50. Valdés Hernández M del C, Ballerini L, Glatz A, Muñoz Maniega S, Gow AJ, Bastin ME, et al. Perivascular spaces in the centrum semiovale at the beginning of the 8th decade of life: effect on cognition and associations with mineral deposition. Brain Imaging Behav. 2019;

51. Ballerini L, Booth T, Valdés Hernández M del C, Wiseman S, Lovreglio R, Muñoz Maniega S, et al. Computational quantification of brain perivascular space morphologies: Associations with vascular risk factors and white matter hyperintensities. A study in the Lothian Birth Cohort 1936. NeuroImage Clin. 2020 Jan 1;25.

52. Desikan RS, Ségonne F, Fischl B, Quinn BT, Dickerson BC, Blacker D, et al. An automated labeling system for subdividing the human cerebral cortex on MRI scans into gyral based regions of interest. Neuroimage. 2006;31(3):968–80.

53. Wright S. Coefficients of Inbreeding and Relationship. Am Nat. 1922 Jul 29;56(645):330–8.

54. Laurie CC, Doheny KF, Mirel DB, Pugh EW, Bierut LJ, Bhangale T, et al. Quality control and quality assurance in genotypic data for genome-wide association studies. Genet Epidemiol [Internet]. 2010 Sep 1 [cited 2020 Feb 10];34(6):591–602. Available from: http://doi.wiley.com/10.1002/gepi.20516

55. Reed E, Nunez S, Kulp D, Qian J, Reilly MP, Foulkes AS. A guide to genome-wide association analysis and post-analytic interrogation. Stat Med [Internet]. 2015 Dec 10 [cited 2020 Feb 7];34(28):3769–92. Available from: http://doi.wiley.com/10.1002/sim.6605

56. Del C. Valdes Hernandez M, Piper RJ, Wang X, Deary IJ, Wardlaw JM. Towards the automatic computational assessment of enlarged perivascular spaces on brain magnetic resonance images: A systematic review. J Magn Reson Imaging. 2013;38(4):774–85.

57. Hilal S, Tan CS, Adams HHH, Habes M, Mok V, Venketasubramanian N, et al. Enlarged perivascular spaces and cognition: A meta-analysis of 5 population-based studies. Neurology [Internet]. 2018 Aug 28 [cited 2019 May 12];91(9):e832.#x2013;42. Available from: http://www.ncbi.nlm.nih.gov/pubmed/30068634

58. Kwee RM, Kwee TC. Virchow-Robin Spaces at MR Imaging. RadioGraphics [Internet]. 2007;27(4):1071–86. Available from: http://pubs.rsna.org/doi/10.1148/rg.274065722

59. Mestre H, Du T, Sweeney AM, Liu G, Samson AJ, Peng W, et al. Cerebrospinal fluid influx drives acute ischemic tissue swelling. Science (80-). 2020 Jan 30;eaax7171.

60. Laveskog A, Wang R, Bronge L, Wahlund LO, Qiu C. Perivascular spaces in old age: Assessment, distribution, and correlation with white matter hyperintensities. Am J Neuroradiol. 2018;39(1):70–6.

61. Berdahl JP, Fleischman D, Zaydlarova J, Stinnett S, Rand Allingham R, Fautsch MP. Body mass index has a linear relationship with cerebrospinal fluid pressure. Vol. 53, Investigative Ophthalmology and Visual Science. The Association for Research in Vision and Ophthalmology; 2012. p. 1422–7.

62. Stapleton PA, James ME, Goodwill AG, Frisbee JC. Obesity and vascular dysfunction. Pathophysiology. 2008 Aug;15(2):79–89.

63. Iliff JJ, Wang M, Zeppenfeld DM, Venkataraman A, Plog BA, Liao Y, et al. Cerebral arterial pulsation drives paravascular CSF-interstitial fluid exchange in the murine brain. J Neurosci [Internet]. 2013 Nov 13 [cited 2017 Sep 25];33(46):18190–9. Available from: http://www.ncbi.nlm.nih.gov/pubmed/24227727

64. Zhu Y-C, Tzourio C, Soumaré A, Mazoyer B, Dufouil C, Chabriat H. Severity of dilated Virchow-Robin spaces is associated with age, blood pressure, and MRI markers of small vessel disease: a population-based study. Stroke [Internet]. 2010 Nov 1 [cited 2018 Jan 9];41(11):2483–90. Available from: http://stroke.ahajournals.org/cgi/doi/10.1161/STROKEAHA.110.591586

65. Siervo M, Ruggiero D, Sorice R, Nutile T, Aversano M, Iafusco M, et al. Body mass index is directly associated with biomarkers of angiogenesis and inflammation in children and adolescents. Nutrition. 2012 Mar 1;28(3):262–6.

66. Santos-Galindo M, Acaz-Fonseca E, Bellini MJ, Garcia-Segura LM. Sex differences in the inflammatory response of primary astrocytes to lipopolysaccharide. Biol Sex Differ [Internet]. 2011 Jul 11 [cited 2020 Mar 19];2(1):7. Available from: http://bsd.biomedcentral.com/articles/10.1186/2042-6410-2-7

67. Song TJ, Park JH, Choi K, Chang Y, Moon J, Kim JH, et al. Moderate-to-severe obstructive sleep apnea is associated with cerebral small vessel disease. Sleep Med. 2017 Feb 1;30:36–42.

68. Lee H, Xie L, Yu M, Kang H, Feng T, Deane R, et al. The effect of body posture on brain glymphatic transport. J Neurosci. 2015 Aug 5;35(31):11034–44.

69. Jelicic M, Bosma H, Ponds RWHM, Van Boxtel MPJ, Houx PJ, Jolles J. Subjective sleep problems in later life as predictors of cognitive decline. Report from the Maastricht Ageing Study (MAAS). Int J Geriatr Psychiatry [Internet]. 2002 Jan 1 [cited 2020 Mar 3];17(1):73–7. Available from: http://doi.wiley.com/10.1002/gps.529

70. Waller KL, Mortensen EL, Avlund K, Osler M, Fagerlund B, Lauritzen M, et al. Subjective sleep quality and daytime sleepiness in late midlife and their association with age-related changes in cognition. Sleep Med. 2016 Jan 1;17:165–73.

71. Ringstad G, Vatnehol SAS, Eide PK. Glymphatic MRI in idiopathic normal pressure hydrocephalus. Brain. 2017;140(10):2691–705.

72. Eide PK, Ringstad G. Delayed clearance of cerebrospinal fluid tracer from entorhinal cortex in idiopathic normal pressure hydrocephalus: A glymphatic magnetic resonance imaging study. J Cereb Blood Flow Metab [Internet]. 2019 Jul [cited 2020 Jan 17];39(7):1355–68. Available from: http://www.ncbi.nlm.nih.gov/pubmed/29485341

73. Xie L, Kang H, Xu Q, Chen MJ, Liao Y, Thiyagarajan M, et al. Sleep drives metabolite clearance from the adult brain. Science (80-). 2013;342(6156):373–7.

74. Sepehrband F, Cabeen RP, Choupan J, Barisano G, Law M, Toga AW. Perivascular space fluid contributes to diffusion tensor imaging changes in white matter. Neuroimage [Internet]. 2019 Apr 30 [cited 2019 May 1];197:243–54. Available from: https://www.sciencedirect.com/science/article/pii/S1053811919303623

75. Thomas C, Sadeghi N, Nayak A, Trefler A, Sarlls J, Baker CI, et al. Impact of time-of-day on diffusivity measures of brain tissue derived from diffusion tensor imaging. Neuroimage. 2018 Jun 1;173:25–34.

76. Dreha-Kulaczewski S, Joseph AA, Merboldt KD, Ludwig HC, Gärtner J, Frahm J. Inspiration is the major regulator of human CSF flow. J Neurosci. 2015 Feb 11;35(6):2485–91.

77. Le KTT, Matzaraki V, Netea MG, Wijmenga C, Moser J, Kumar V. Functional annotation of genetic loci associated with sepsis prioritizes immune and endothelial cell pathways. Front Immunol. 2019;10(AUG).

78. Erlich SS, McComb JG, Hyman S, Weiss MH. Ultrastructural morphology of the olfactory pathway for cerebrospinal fluid drainage in the rabbit. J Neurosurg [Internet]. 1986 Mar [cited 2020 Mar 4];64(3):466–73. Available from: http://www.ncbi.nlm.nih.gov/pubmed/3950724

79. Löwhagen P, Johansson BB, Nordborg C. The nasal route of cerebrospinal fluid drainage in man. A light–microscope study. Neuropathol Appl Neurobiol [Internet]. 1994 Dec 1 [cited 2020 Mar 4];20(6):543–50. Available from: http://doi.wiley.com/10.1111/j.1365-2990.1994.tb01008.x

80. Koh L, Zakharov A, Nagra G, Armstrong D, Friendship R, Johnston M. Development of cerebrospinal fluid absorption sites in the pig and rat: Connections between the subarachnoid space and lymphatic vessels in the olfactory turbinates. Anat Embryol (Berl). 2006 Aug 10;211(4):335–44.

81. Kim HJ, Kim PK, Bae SM, Son HN, Thoudam DS, Kim JE, et al. Transforming growth factor-β-induced protein (TGFBIp/β ig-h3) activates platelets and promotes thrombogenesis. Blood. 2009;114(25):5206–15.

82. Mestre H, Kostrikov S, Mehta RI, Nedergaard M. Perivascular spaces, glymphatic dysfunction, and small vessel disease. Clin Sci [Internet]. 2017;131(17):2257–74. Available from: http://clinsci.org/lookup/doi/10.1042/CS20160381

