## Supplementary tables for "Body mass index, time of day, and genetics affect perivascular spaces in the white matter"

### **Supplementary material**

### **Supplementary Figure Legends**

**Supplementary figure 1.** Scatterplots showing the relationship between the difference in body mass index (BMI) and the difference in perivascular space (PVS) in each couple of monozygotic twins (**a**), dizygotic twins (**b**), non-twin siblings (**c**). Spearman’s rank correlation coefficient.

**Supplementary figure 2.** Scatterplots showing the relationship of years of education with body mass index (BMI) (**a**) and perivascular space (PVS) ratio (**b**). Spearman’s rank correlation coefficient.

### **Supplementary Tables**

**Supplementary Table 1.** Inclusion and exclusion criteria for the participants enrolled in the Human Connectome Project (S900 release)(33).

**Inclusion criteria**

- Age between 22-37

- No prior history of psychiatric disorders, substance abuse, neurological or cardiovascular diseases, as indicated by no report of medical diagnosis, no hospitalization, and no pharmacologic or behavioral treatment.

**Exclusion criteria**

- Any genetic disorder

- Current use of chemotherapy or immunomodulatory agents

- History of radiation or chemotherapy that could affect the brain

- Sickle cell disease

- Thyroid hormone treatment within 12 months before the enrollment

- Treatment for diabetes

- Head injury followed by neurological symptoms, such as loss of consciousness for more than 30 minutes or amnesia or change in mental status for more than 24 hours, and/or CT findings consistent with traumatic brain injury.

- Ineligible to undergo an MRI scan: pregnancy, metal or devices in the body not compatible with MRI (e.g., cardiac pacemaker, cochlear implant, aneurism clip), and/or suffering moderate to severe claustrophobia were reasons for being excluded from the study.

- Non-twins born prior to 37 weeks of gestation and twins born prior to 34 weeks of gestation have been excluded since preterm birth has been shown to perturb the development of the brain.(34)

**Supplementary Table 2.** Perivascular space ratio in each region of interest (ROI), obtained by dividing the perivascular space volume computed in the ROI by the white matter volume of the same ROI. For each ROI, the first row represents the sum of the PVS ratio computed in the two hemispheres, while the PVS ratio of each side is reported in the second and third row (right and left side, respectively). Data are mean ± standard deviation (range: minumum-maximum value). The adjusted p-values refer to the Wilcoxon matched-pairs signed rank test performed to compare the PVS ratio on the right side of each ROI with the corresponding contralateral side. The adjusted p-values that are not significant after controlling for the false discovery rate are reported in red.

| **ROI** | **Side** | **Male (n=394)** | **Female (n=503)** | **Overall (n=897)** | **Adjusted p-value** |
| --- | --- | --- | --- | --- | --- |
| Bankssts |  | 2.11 ± 1.08 (0.24-7.13) | 2.06 ± 0.96 (0.29-5.71) | 2.08 ± 1.01 (0.24-7.13) |  |
|  | Right | 1.17 ± 0.67 (0.02-4.61) | 1.14 ± 0.58 (0.05-3.42) | 1.15 ± 0.62 (0.02-4.61) | 2.50E-30 |
|  | Left | 0.94 ± 0.52 (0-3.19) | 0.92 ± 0.53 (0-2.72) | 0.93 ± 0.53 (0-3.19) |  |
| Caudalanteriorcingulate | | 3.55 ± 2.02 (0.28-9.96) | 2.74 ± 1.56 (0.17-9.98) | 3.09 ± 1.82 (0.17-9.98) |  |
|  | Right | 1.7 ± 1.08 (0.14-5.14) | 1.31 ± 0.83 (0.03-5.42) | 1.48 ± 0.96 (0.03-5.42) | 2.00E-11 |
|  | Left | 1.85 ± 1.03 (0.13-5.31) | 1.43 ± 0.84 (0.05-4.86) | 1.61 ± 0.95 (0.05-5.31) |  |
| Caudalmiddlefrontal | | 2.39 ± 1.26 (0.18-6.75) | 2.08 ± 1.07 (0.29-6.15) | 2.22 ± 1.17 (0.18-6.75) |  |
|  | Right | 1.05 ± 0.63 (0.08-3.33) | 0.93 ± 0.52 (0.07-2.82) | 0.98 ± 0.57 (0.07-3.33) | 1.60E-70 |
|  | Left | 1.34 ± 0.69 (0.05-3.67) | 1.15 ± 0.6 (0.09-3.33) | 1.24 ± 0.65 (0.05-3.67) |  |
| Cuneus |  | 0.55 ± 0.54 (0-4.81) | 0.46 ± 0.46 (0-2.77) | 0.5 ± 0.5 (0-4.81) |  |
|  | Right | 0.29 ± 0.31 (0-3.17) | 0.23 ± 0.27 (0-1.62) | 0.26 ± 0.29 (0-3.17) | 0.24 |
|  | Left | 0.26 ± 0.29 (0-1.82) | 0.22 ± 0.24 (0-1.2) | 0.24 ± 0.26 (0-1.82) |  |
| Entorhinal |  | 0.5 ± 0.6 (0-3.48) | 0.24 ± 0.33 (0-2.2) | 0.35 ± 0.48 (0-3.48) |  |
|  | Right | 0.15 ± 0.26 (0-1.86) | 0.08 ± 0.16 (0-1.39) | 0.11 ± 0.21 (0-1.86) | 2.40E-30 |
|  | Left | 0.34 ± 0.45 (0-2.91) | 0.16 ± 0.27 (0-2.06) | 0.24 ± 0.37 (0-2.91) |  |
| Fusiform |  | 1.56 ± 0.81 (0.12-5.16) | 1.46 ± 0.78 (0.12-6.88) | 1.51 ± 0.8 (0.12-6.88) |  |
|  | Right | 0.8 ± 0.45 (0.06-2.72) | 0.77 ± 0.44 (0.02-3.76) | 0.78 ± 0.44 (0.02-3.76) | 6.40E-08 |
|  | Left | 0.76 ± 0.43 (0.05-2.91) | 0.7 ± 0.41 (0.03-3.13) | 0.72 ± 0.42 (0.03-3.13) |  |
| Inferiorparietal | | 2.67 ± 1.29 (0.35-8.01) | 2.58 ± 1.23 (0.26-7.29) | 2.62 ± 1.25 (0.26-8.01) |  |
|  | Right | 1.47 ± 0.7 (0.18-4.33) | 1.4 ± 0.68 (0.2-4.07) | 1.43 ± 0.69 (0.18-4.33) | 2.00E-64 |
|  | Left | 1.2 ± 0.63 (0.14-4.39) | 1.18 ± 0.6 (0.07-3.56) | 1.19 ± 0.61 (0.07-4.39) |  |
| Inferiortemporal | | 1.48 ± 0.83 (0.11-5.22) | 1.39 ± 0.77 (0.08-5.01) | 1.43 ± 0.8 (0.08-5.22) |  |
|  | Right | 0.69 ± 0.43 (0.01-2.83) | 0.68 ± 0.4 (0.01-2.53) | 0.69 ± 0.41 (0.01-2.83) | 2.30E-07 |
|  | Left | 0.78 ± 0.45 (0.01-2.74) | 0.71 ± 0.43 (0-2.69) | 0.74 ± 0.44 (0-2.74) |  |
| Isthmuscingulate | | 3.63 ± 1.77 (0.39-10.15) | 2.71 ± 1.27 (0.24-8.12) | 3.11 ± 1.58 (0.24-10.15) |  |
|  | Right | 1.96 ± 0.99 (0.25-5.31) | 1.4 ± 0.7 (0.07-4.84) | 1.65 ± 0.89 (0.07-5.31) | 5.00E-30 |
|  | Left | 1.67 ± 0.83 (0.12-4.84) | 1.3 ± 0.63 (0.12-3.77) | 1.47 ± 0.75 (0.12-4.84) |  |
| Lateraloccipital | | 1.06 ± 0.73 (0.03-4.61) | 1 ± 0.64 (0.03-3.89) | 1.03 ± 0.68 (0.03-4.61) |  |
|  | Right | 0.59 ± 0.41 (0-2.33) | 0.56 ± 0.37 (0.02-2.4) | 0.57 ± 0.39 (0-2.4) | 3.40E-49 |
|  | Left | 0.47 ± 0.36 (0.02-2.28) | 0.44 ± 0.3 (0-1.63) | 0.45 ± 0.33 (0-2.28) |  |
| Lateralorbitofrontal | | 2.54 ± 1.44 (0.39-8.15) | 2.26 ± 1.27 (0.32-8.67) | 2.38 ± 1.36 (0.32-8.67) |  |
|  | Right | 1.19 ± 0.71 (0.15-4.23) | 1.11 ± 0.64 (0.16-4.59) | 1.14 ± 0.67 (0.15-4.59) | 1.60E-08 |
|  | Left | 1.35 ± 0.8 (0.14-4.31) | 1.15 ± 0.7 (0.12-5.29) | 1.24 ± 0.75 (0.12-5.29) |  |
| Lingual |  | 1.06 ± 0.6 (0.07-4.29) | 1 ± 0.57 (0-3.52) | 1.03 ± 0.59 (0-4.29) |  |
|  | Right | 0.64 ± 0.38 (0.02-2.36) | 0.6 ± 0.36 (0-2.28) | 0.62 ± 0.37 (0-2.36) | 3.50E-69 |
|  | Left | 0.42 ± 0.32 (0-2.09) | 0.4 ± 0.3 (0-1.98) | 0.41 ± 0.31 (0-2.09) |  |
| Medialorbitofrontal | | 1.46 ± 1.09 (0.05-7.15) | 1.24 ± 0.88 (0.08-6.16) | 1.33 ± 0.98 (0.05-7.15) |  |
|  | Right | 0.64 ± 0.61 (0-3.89) | 0.55 ± 0.51 (0-3.7) | 0.59 ± 0.56 (0-3.89) | 1.90E-41 |
|  | Left | 0.81 ± 0.54 (0-3.42) | 0.69 ± 0.45 (0.03-2.78) | 0.74 ± 0.49 (0-3.42) |  |
| Middletemporal | | 1.51 ± 0.81 (0.11-4.78) | 1.43 ± 0.76 (0.11-4.62) | 1.46 ± 0.78 (0.11-4.78) |  |
|  | Right | 0.73 ± 0.43 (0.02-2.7) | 0.7 ± 0.4 (0.06-2.73) | 0.71 ± 0.42 (0.02-2.73) | 0.001 |
|  | Left | 0.79 ± 0.44 (0.07-2.84) | 0.73 ± 0.42 (0.02-2.66) | 0.75 ± 0.43 (0.02-2.84) |  |
| Parahippocampal | | 0.83 ± 0.62 (0-4.12) | 0.58 ± 0.56 (0-3.76) | 0.69 ± 0.6 (0-4.12) |  |
|  | Right | 0.37 ± 0.32 (0-1.98) | 0.28 ± 0.35 (0-2.86) | 0.32 ± 0.34 (0-2.86) | 0.005 |
|  | Left | 0.45 ± 0.41 (0-3) | 0.3 ± 0.35 (0-2.73) | 0.36 ± 0.38 (0-3) |  |
| Paracentral |  | 1.42 ± 0.86 (0.04-5.19) | 1.24 ± 0.77 (0.03-4.41) | 1.32 ± 0.82 (0.03-5.19) |  |
|  | Right | 0.8 ± 0.5 (0.02-3.21) | 0.68 ± 0.44 (0.03-2.55) | 0.73 ± 0.47 (0.02-3.21) | 3.80E-35 |
|  | Left | 0.62 ± 0.42 (0-2.09) | 0.56 ± 0.39 (0-2.34) | 0.59 ± 0.4 (0-2.34) |  |
| Parsopercularis | | 1.96 ± 1.26 (0.03-7.17) | 1.62 ± 1.11 (0.06-7.45) | 1.77 ± 1.19 (0.03-7.45) |  |
|  | Right | 0.99 ± 0.69 (0-3.75) | 0.84 ± 0.6 (0-3.18) | 0.91 ± 0.65 (0-3.75) | 0.034 |
|  | Left | 0.97 ± 0.65 (0-3.43) | 0.77 ± 0.59 (0-4.28) | 0.86 ± 0.63 (0-4.28) |  |
| Parsorbitalis | | 0.85 ± 0.76 (0-3.9) | 0.78 ± 0.71 (0-4) | 0.81 ± 0.73 (0-4) |  |
|  | Right | 0.41 ± 0.42 (0-2.37) | 0.37 ± 0.36 (0-1.89) | 0.39 ± 0.39 (0-2.37) | 0.28 |
|  | Left | 0.44 ± 0.44 (0-2.25) | 0.41 ± 0.46 (0-2.87) | 0.42 ± 0.45 (0-2.87) |  |
| Parstriangularis | | 1.5 ± 1.08 (0-5.84) | 1.28 ± 0.86 (0-4.81) | 1.37 ± 0.96 (0-5.84) |  |
|  | Right | 0.64 ± 0.53 (0-2.85) | 0.57 ± 0.44 (0-2.94) | 0.6 ± 0.48 (0-2.94) | 2.00E-36 |
|  | Left | 0.86 ± 0.62 (0-3.21) | 0.71 ± 0.5 (0-3.2) | 0.78 ± 0.56 (0-3.21) |  |
| Pericalcarine | | 2.1 ± 1.28 (0-6.57) | 2.14 ± 1.26 (0-7.37) | 2.12 ± 1.27 (0-7.37) |  |
|  | Right | 1.03 ± 0.69 (0-3.45) | 1.05 ± 0.7 (0-3.42) | 1.04 ± 0.69 (0-3.45) | 1 |
|  | Left | 1.07 ± 0.86 (0-4.81) | 1.09 ± 0.88 (0-4.65) | 1.08 ± 0.87 (0-4.81) |  |
| Postcentral |  | 1.61 ± 0.96 (0.06-5.82) | 1.4 ± 0.82 (0.06-4.88) | 1.49 ± 0.89 (0.06-5.82) |  |
|  | Right | 0.79 ± 0.49 (0.02-3.07) | 0.69 ± 0.42 (0.02-2.53) | 0.73 ± 0.45 (0.02-3.07) | 0.017 |
|  | Left | 0.83 ± 0.53 (0.03-3.56) | 0.71 ± 0.45 (0-2.6) | 0.76 ± 0.49 (0-3.56) |  |
| Posteriorcingulate | | 2.6 ± 1.32 (0.33-6.97) | 2.11 ± 1.07 (0.34-6.47) | 2.33 ± 1.21 (0.33-6.97) |  |
|  | Right | 1.35 ± 0.71 (0.15-3.77) | 1.08 ± 0.58 (0.15-3.36) | 1.2 ± 0.65 (0.15-3.77) | 3.30E-06 |
|  | Left | 1.25 ± 0.65 (0.12-3.61) | 1.03 ± 0.55 (0.1-3.46) | 1.13 ± 0.61 (0.1-3.61) |  |
| Precentral |  | 2.23 ± 1.2 (0.23-7.12) | 1.98 ± 1.06 (0.25-6.68) | 2.09 ± 1.13 (0.23-7.12) |  |
|  | Right | 1.11 ± 0.62 (0.12-3.8) | 0.98 ± 0.53 (0.06-3.06) | 1.04 ± 0.58 (0.06-3.8) | 0.57 |
|  | Left | 1.11 ± 0.61 (0.11-3.78) | 1 ± 0.55 (0.05-3.62) | 1.05 ± 0.58 (0.05-3.78) |  |
| Precuneus |  | 2.65 ± 1.11 (0.37-7.24) | 2.45 ± 1.06 (0.52-6.35) | 2.54 ± 1.09 (0.37-7.24) |  |
|  | Right | 1.47 ± 0.61 (0.24-3.85) | 1.33 ± 0.59 (0.28-3.69) | 1.39 ± 0.61 (0.24-3.85) | 5.10E-74 |
|  | Left | 1.18 ± 0.55 (0.13-3.39) | 1.11 ± 0.52 (0.22-3.3) | 1.14 ± 0.53 (0.13-3.39) |  |
| Rostralanteriorcingulate | | 3.15 ± 2.05 (0.3-13.05) | 2.53 ± 1.61 (0.15-11.6) | 2.8 ± 1.84 (0.15-13.05) |  |
|  | Right | 1.57 ± 1.1 (0.07-6.04) | 1.25 ± 0.86 (0-5.88) | 1.39 ± 0.99 (0-6.04) | 0.57 |
|  | Left | 1.58 ± 1.05 (0.09-7.26) | 1.28 ± 0.84 (0-5.72) | 1.41 ± 0.95 (0-7.26) |  |
| Rostralmiddlefrontal | | 1.97 ± 1.26 (0.09-7.52) | 1.81 ± 1.06 (0.15-6.13) | 1.88 ± 1.16 (0.09-7.52) |  |
|  | Right | 0.84 ± 0.59 (0.03-3.67) | 0.8 ± 0.51 (0.04-3.39) | 0.82 ± 0.55 (0.03-3.67) | 2.50E-96 |
|  | Left | 1.13 ± 0.7 (0.02-4.28) | 1.01 ± 0.59 (0.09-4.11) | 1.06 ± 0.64 (0.02-4.28) |  |
| Superiorfrontal | | 2.49 ± 1.2 (0.33-7.13) | 2.22 ± 1.09 (0.25-6.55) | 2.33 ± 1.15 (0.25-7.13) |  |
|  | Right | 1.21 ± 0.61 (0.13-3.45) | 1.09 ± 0.55 (0.1-3.59) | 1.14 ± 0.58 (0.1-3.59) | 3.40E-09 |
|  | Left | 1.27 ± 0.61 (0.13-3.73) | 1.13 ± 0.57 (0.12-3.3) | 1.19 ± 0.59 (0.12-3.73) |  |
| Superiorparietal | | 2.62 ± 1.22 (0.52-6.31) | 2.41 ± 1.07 (0.45-6) | 2.5 ± 1.14 (0.45-6.31) |  |
|  | Right | 1.33 ± 0.63 (0.23-3.54) | 1.23 ± 0.56 (0.2-3.27) | 1.27 ± 0.6 (0.2-3.54) | 2.10E-05 |
|  | Left | 1.29 ± 0.62 (0.28-3.4) | 1.18 ± 0.54 (0.13-2.9) | 1.23 ± 0.58 (0.13-3.4) |  |
| Superiortemporal | | 1.44 ± 0.81 (0.11-4.2) | 1.26 ± 0.74 (0.06-4.52) | 1.34 ± 0.78 (0.06-4.52) |  |
|  | Right | 0.59 ± 0.4 (0.02-2.45) | 0.54 ± 0.38 (0.01-2.49) | 0.57 ± 0.39 (0.01-2.49) | 3.80E-68 |
|  | Left | 0.85 ± 0.47 (0.01-2.73) | 0.71 ± 0.43 (0.02-2.46) | 0.77 ± 0.45 (0.01-2.73) |  |
| Supramarginal | | 3.25 ± 1.58 (0.25-8.69) | 2.9 ± 1.37 (0.44-7.75) | 3.06 ± 1.48 (0.25-8.69) |  |
|  | Right | 1.6 ± 0.8 (0.14-4.35) | 1.4 ± 0.68 (0.1-3.72) | 1.49 ± 0.74 (0.1-4.35) | 3.10E-07 |
|  | Left | 1.66 ± 0.85 (0.03-4.79) | 1.5 ± 0.76 (0.19-4.16) | 1.57 ± 0.8 (0.03-4.79) |  |
| Frontalpole |  | 0.32 ± 0.49 (0-2.96) | 0.24 ± 0.4 (0-2.77) | 0.28 ± 0.44 (0-2.96) |  |
|  | Right | 0.17 ± 0.32 (0-1.83) | 0.13 ± 0.27 (0-1.68) | 0.15 ± 0.29 (0-1.83) | 0.063 |
|  | Left | 0.15 ± 0.32 (0-2.4) | 0.11 ± 0.27 (0-2.77) | 0.13 ± 0.3 (0-2.77) |  |
| Temporalpole | | 0.89 ± 0.86 (0-4.32) | 0.62 ± 0.71 (0-6.45) | 0.74 ± 0.79 (0-6.45) |  |
|  | Right | 0.36 ± 0.45 (0-3.43) | 0.27 ± 0.38 (0-2.78) | 0.31 ± 0.41 (0-3.43) | 2.10E-12 |
|  | Left | 0.53 ± 0.56 (0-3.19) | 0.35 ± 0.47 (0-3.67) | 0.43 ± 0.52 (0-3.67) |  |
| Transversetemporal | | 1.52 ± 1.26 (0-7.15) | 1.7 ± 1.37 (0-8.34) | 1.62 ± 1.33 (0-8.34) |  |
|  | Right | 0.74 ± 0.81 (0-5.09) | 0.88 ± 0.87 (0-4.87) | 0.82 ± 0.85 (0-5.09) | 1 |
|  | Left | 0.78 ± 0.72 (0-3.76) | 0.82 ± 0.78 (0-4.88) | 0.81 ± 0.75 (0-4.88) |  |
| Insula |  | 3.25 ± 1.12 (0.82-7.76) | 2.87 ± 0.95 (0.96-7.43) | 3.04 ± 1.05 (0.82-7.76) |  |
|  | Right | 1.58 ± 0.6 (0.41-4.52) | 1.43 ± 0.52 (0.33-3.87) | 1.5 ± 0.56 (0.33-4.52) | 0.0013 |
|  | Left | 1.67 ± 0.58 (0.41-3.92) | 1.44 ± 0.49 (0.38-3.56) | 1.54 ± 0.55 (0.38-3.92) |  |

**Supplementary Table 3.** Results of the ANCOVA model testing the effects of gender and BMI on PVS ratio after controlling for age. BMI groups: 1. BMI < 20; 2. BMI between 20 and 25; 3. BMI between 25 and 30; 4. BMI > 30. Significant p-values after controlling for the false discovery rate are marked with *.

|  | *F*-test statistic | p-value | η^2^ |
| --- | --- | --- | --- |
| Age | 22.209 | 2.85E-06* | 0.025 |
| Gender | 42.572 | 1.16E-10* | 0.047 |
| BMI groups | 15.552 | 7.34E-10* | 0.051 |
| Gender:BMI groups interaction | 2.698 | 0.045 | 0.009 |
